# Novel cell wall-associated genes that enable *Cryptococcus neoformans* to evade dectin-1-mediated innate immune recognition

**DOI:** 10.64898/2026.07.27.740439

**Authors:** Keigo Ueno, Akiko Nagamori, Nahoko Oniyama Honkyu, Daisuke Yamanaka, Ken Miyazawa, Ami Koizumi, Kyung J. Kwon-Chung, Yoshitsugu Miyazaki

## Abstract

The fungal pathogen *Cryptococcus neoformans* contains about 200 µg of β-1,3-glucan (1,3BG) per 1 mg of dry cell weight when grown under standard culture conditions (YPD medium at 30 °C under aerobic conditions). However, 1,3BG exposure is tightly suppressed, even in capsule-deficient strains, allowing the fungus to evade recognition by the immune receptor dectin-1 and anti-1,3BG antibodies. Although other pathogenic fungi mask 1,3BG with α-1,3-glucan (1,3AG) to evade dectin-1 recognition, the factors responsible for 1,3BG masking and dectin-1 evasion in *C. neoformans* remain incompletely understood. To identify capsule-independent 1,3BG masking and dectin-1-evasion factors, we generated a series of cell wall-related gene deletion strains in the capsule-deficient strain *cap59*Δ by using CRISPR-Cas9 and screened for mutants that failed to evade dectin-1 binding. We found eight deletants (*cap59*Δ/*mpk1*Δ, *cap59*Δ/*chs3*Δ, *cap59*Δ/*kre5*Δ, *cap59*Δ/*crz1*Δ, *cap59*Δ/*kre6*Δ/*skn1*Δ, *cap59*Δ/*hxl1*Δ, *cap59*Δ/*uge1*Δ, and *cap59*Δ/*ugt1*Δ) that exhibited increased binding to dectin-1 or anti-1,3BG antibody, or both. As a similar phenotype was not observed in *cap59*Δ/*ags1*Δ, 1,3AG-mediated masking of 1,3BG appears to play a limited role in *C. neoformans*. These eight deletants induced significantly greater secretion of IL-6 and IL-1β from dendritic cells (DCs) than did *cap59*Δ. This enhanced inflammatory response was markedly attenuated in dectin-1-deficient DCs, indicating that the increased immunogenicity was driven by 1,3BG exposure and subsequent dectin-1 recognition. Collectively, these findings demonstrate that multiple genes involved in maintaining cell wall integrity—including those involved in β-1,6-glucan and chitosan biosynthesis—are essential for regulating 1,3BG exposure and enabling *C. neoformans* to evade dectin-1-mediated immune recognition.

**Highlights:**

- We identified new capsule-independent β-1,3-glucan-masking genes in *C. neoformans*
- Gene deletants had higher dectin-1 deposition than parental acapsular *cap59*Δ
- Deletion of these genes enhanced IL-6 and IL-1β secretion by dendritic cells
- The enhanced cytokine response was suppressed in dendritic cells lacking dectin-1.
- Deletant strains may serve as new whole-cell antigens for cryptococcal vaccines

## 1. Introduction

*Cryptococcus neoformans* is a fungal pathogen that causes cryptococcosis primarily in immunosuppressed patients. However, cryptococcosis can also occur in apparently immunocompetent individuals, suggesting that *C. neoformans* possesses a remarkable ability to evade the host immune system (Denham and Brown, 2018; Diniz-Lima et al., 2022).

Capsule polysaccharides, particularly glucuronoxylomannan (GXM), are the major determinants of immune evasion, as they mask fungal antigens and prevent recognition by the innate immune receptor CD11b (Ueno et al., 2021). Capsule-deficient strains are avirulent and are rapidly eliminated from infected lungs (Chang and Kwon-Chung, 1994; Ueno et al., 2021). In addition, *C. neoformans* possesses a capsule-independent mechanism that regulates the exposure of β-1,3-glucan (1,3BG), thereby preventing recognition by the innate immune receptor dectin-1 (Bloom et al., 2021, 2019; Farkas et al., 2009; Ueno et al., 2019; Upadhya et al., 2023). The glucan layer of the *C. neoformans* cell wall is composed primarily of β-1,6-glucan (1,6BG) and α-1,3-glucan (1,3AG), but the amount of 1,3BG is lower than that in many other pathogenic fungi (Ankur et al., 2025; Garcia-Rubio et al., 2020; James et al., 1990). Nevertheless, the alkali-insoluble cell wall fraction contains approximately 200 µg of 1,3BG per 1 mg of dry fungal weight, indicating that 1,3BG is present in substantial amounts (Reese et al., 2007). Interestingly, when *C. neoformans* is grown aerobically in standard yeast extract peptone dextrose (YPD) medium at 30 °C, 1,3BG is minimally exposed, and neither dectin-1 nor anti-1,3BG antibodies bind to the cell surface even in acapsular strains (Ueno et al., 2019; Upadhya et al., 2023; Walsh et al., 2017). In contrast, cells grown in synthetic glucose medium—SD [synthetic dextrose] medium or YNB [yeast nitrogen base] glucose medium—or Sabouraud medium expose their 1,3BG irrespective of capsule formation, allowing binding by both dectin-1 and anti-1,3BG antibodies. These cells exhibit enhanced immunogenicity and induce higher levels of inflammatory cytokines in dendritic cells and murine lungs (Ueno et al., 2019; Upadhya et al., 2023). Acidification of SD medium during culture has been identified as one of the factors promoting 1,3BG exposure and increasing immunoregulatory activity (Farkas et al., 2009; Rachini et al., 2007; Ueno et al., 2019; Upadhya et al., 2023). Furthermore, 1,3BG is generally exposed on the surface of the *C. neoformans* spore (Walsh et al., 2017). Collectively, these findings demonstrate that *Cryptococcus* spp. tightly regulate their 1,3BG exposure in response to environmental cues to evade dectin-1-mediated immune recognition. Previous studies have shown that minimization of 1,3BG exposure in *C. neoformans* under thermal stress (YPD, 37 °C, aerobic conditions) involves *CCR4* (Carbon Catabolite Repression 4)-mediated mRNA degradation and transcriptional reprogramming (Bloom et al., 2019). However, the genetic networks controlling 1,3BG exposure under standard culture conditions (YPD, 30 °C, aerobic conditions) remain largely unknown. In *Candida albicans,* 1,3BG is masked during host colonization in response to lactic acid or hypoxic conditions (Ballou et al., 2016; Pradhan et al., 2018), whereas *Aspergillus fumigatus* and *Histoplasma capsulatum* conceal their 1,3BG beneath an α-1,3-glucan (1,3AG) layer to avoid dectin-1 recognition (Beauvais et al., 2013; Rappleye et al., 2007). Whether similar mechanisms operate in *Cryptococcus* spp remains unclear. Because cryptococcal cells cultured aerobically in YPD at 30 °C are used widely in infection models and host–pathogen interaction studies, elucidation of the mechanisms regulating 1,3BG exposure and dectin-1 evasion under these standard conditions will facilitate experimental design and improve our understanding of cryptococcal immune evasion.

Several cryptococcal mutants with impaired immune evasion have been reported, including *mar1*Δ (Esher et al., 2018), *rim101*Δ(O’Meara et al., 2013), *chs3*Δ (Hole et al., 2020), *pdr6*Δ (Winski et al., 2025), and *MAY1* overexpression strain (*MAY1*^oe^) (Li et al., 2024). These highly immunogenic strains induce excessive inflammatory responses in dendritic cells, macrophages, and the lungs of infected mice and, in some cases, lead to fatal outcomes (Hole et al., 2020; O’Meara et al., 2013). Although the cell wall properties of these strains have been characterized, none has been reported to expose its 1,3BG. Highly immunogenic cryptococcal strains, such as *cap59*Δ grown in SD medium, have been used successfully as whole-cell vaccine antigens, and the increasing number of such studies highlights the importance of developing additional highly immunogenic strains of *Cryptococcus* for vaccine development (Ueno et al., 2023b, 2025).

Our objectives here were to: (1) construct new *C. neoformans* strains with enhanced adjuvant activity; (2) characterize the binding properties of innate immune receptors, particularly dectin-1, as well as lectins to these strains; and (3) determine whether heightened immunostimulatory activity is mediated by dectin-1. To achieve these aims, we used the CRISPR-Cas9 method to generate a series of cell wall-related gene-knockout mutants, with *C. neoformans cap59*Δ as the parental strain. From these mutants, we identified highly immunogenic strains that exposed their 1,3BG and were recognized in a dectin-1-dependent manner. These findings reveal a previously unrecognized capsule-independent mechanism of innate immune evasion and provide novel whole-cell vaccine candidates for the development of cryptococcal vaccines.

## 2. Materials and Methods

### 2.1. Ethics

All animal experiments were conducted in accordance with the guidelines for Animal Experiments of the National Institute of Infectious Diseases, Japan (approval number 126004), and complied with ethical principles and institutional policies.

### 2.2. Mice

C57BL/6J mice were purchased from Japan SLC Inc. Dectin-1 knockout (KO) mice were kindly provided by Dr. Shinobu Saijo (Saijo et al., 2007). Mice were maintained in a specific-pathogen-free environment at the animal experiment facility of the National Institute of Infectious Diseases of Japan.

### 2.3. Construction of cryptococcal gene-knockout strains by using CRISPR-Cas9

The target genes are listed in Table 1, and the strains investigated are listed in Table 2. Genetic manipulation of cryptococcal cells was performed by using a CRISPR-Cas9 system (Fig. S1)(Huang et al., 2021). The plasmids for this system were obtained from a nonprofit repository, Addgene (https://www.addgene.org). In this approach, three linear DNA fragments amplified by PCR were introduced into *C. neoformans* by using electroporation, as described below (Fig. S1A). These fragments encoded (1) Cas9, (2) a guide RNA (gRNA) expression cassette containing gRNA and scaffold, and (3) a selectable marker containing resistance to neomycin (*NEO)*, hygromycin B (*HYG),* or nourseothricin (*NAT)*. When the linear DNA contains an appropriate promoter and terminator, the target protein is transiently expressed from the introduced fragment (Huang et al., 2021). Primers used in this study are listed in Table S1.

**Table 1.**
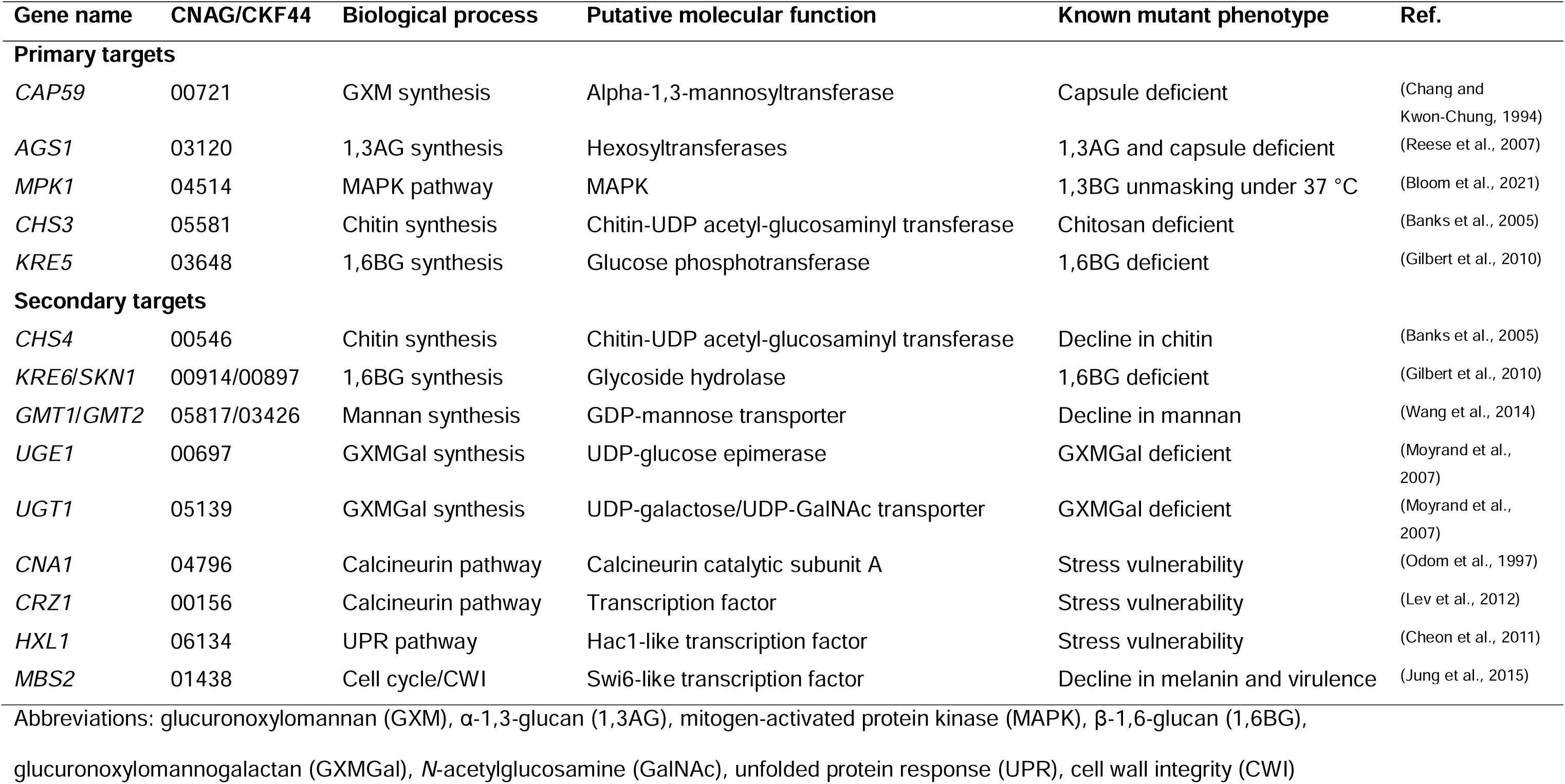
Cell wall-related genes analyzed.

**Table 2.**
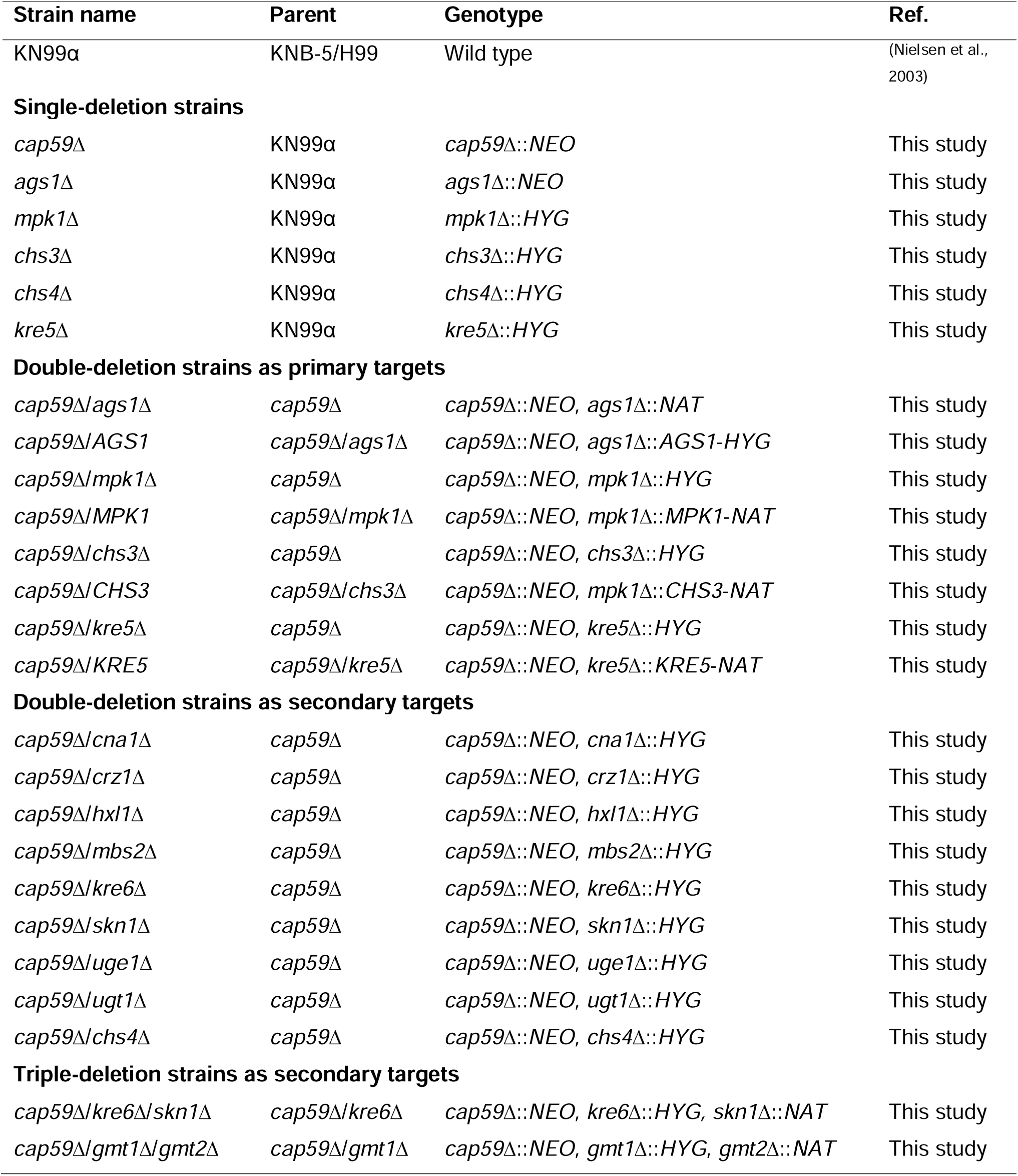
Strains.

The Cas9 fragment was amplified from plasmid pBMH2403 (Fig. S1A) by using primers M13_F and M13_R. Cas9 is responsible for cleaving target DNA; to instruct the cleavage site, gRNA-expressing DNA is also required. In preliminary experiments, cleavage at a single site in the target gene resulted in low recombination efficiency; therefore, we adopted a strategy of using two guide sequences for a single target gene to cleave at two sites—one upstream and one downstream of the target gene (Fig. S1B). To select a guide sequence, the open reading frame (ORF) sequence of *YFG* (your favorite genes) was submitted to the website [http://grna.ctegd.uga.edu (RNA-guided nuclease selection = SpCas9, Genome = *C. neoformans* H99 FungiDB-26). The sequences with the highest possible score were selected. This guide sequence was combined with the scaffold region and is transcribed by the CnU6 promoter (Fig. S1A, Red). This fragment was amplified by overlap-extension PCR using six primers (Fig. S1A, Red). As an example, for cleavage upstream of the MPK1 ORF, plasmid pBHM2329 was used as a template to amplify two types of DNA fragments by using primers M13_F and 4514_g105-R for one, and 4514_g105-F and M13_R for the other. Primers 4514_g105-F and -R contained the guide sequence 5D-GTAGCATTCTTCAACAGCCA-3D (Fig. S1A, Red). These two types of fragments were used as templates to amplify a single DNA fragment by using primers P33 and P34_KU (Fig. S1A, left and right black arrows); this was the DNA fragment for transcribing the guide RNA. Similarly, the gRNA-expressing fragment was amplified for the downstream region by using the same method, with the primer combination changed.

The disruption cassettes contained *NEO*, *NAT*, or *HYG* selectable markers. These are regulated by the *ACT1* promoter and the *TRP1* terminator (Fig. S1A, S1B). Plasmids pJAF1, pJAF13, and pJAF15 were used as templates for the cassette amplification (Fraser et al., 2003). The *MPK1* disruption cassette was amplified from plasmid pJAF15 by using primers 4514-disF1 and 4514-disR1, which include a 56-bp homologous sequence required for the recombination that matches the 56-bp sequences upstream and downstream of the cleavage site (Fig. S1A, S1B: green bars).

To restore the target genes in the deletion mutants, the target gene locus was cloned into plasmid pJAF13 or pJAF15 and then used to amplify the complement cassette (Fig. S1A, far right). *AGS1* and *MPK1*/*CHS3*/*KRE5* were replaced by the *NAT* and *HYG* markers, respectively; thus, *AGS1* and *MPK1*/*CHS3*/*KRE5* loci were cloned into pJAF15 (*HYG*) and pJAF13 (*NAT*), respectively. For this cloning procedure, two amplified DNA fragments were ligated using In-Fusion^®^ Snap Assembly Master Mix (Takara Bio). Vector fragments were amplified from plasmids pJAF13 or pJAF15 with the primers Univ-compmaking_F and Univ-compmaking_R. Insert fragments were amplified from genomic DNA of the *C. neoformans* KN99 strain by using primers mpk1_compF1 and mpk1_compR1. The cloned nucleotide sequences were verified by sequencing with the primers M13_R, UnivCheck-R, mpk1_sc710, mpk1_sc1416, mpk1_sc2096, mpk1_sc2598, and mpk1_sc3189, and they were confirmed to match the database reference sequences. The complement cassette was amplified from a plasmid carrying the target gene by using primers mpk1 cmpcst F1 and 1562-compR3. Primer 1562-compR3 is a universal primer that anneals upstream of the *ACT1* promoter in pJAF13/15. For the double cleavage and recombination at the *NAT* or *HYG* locus (Fig. S1C), gRNA-expression fragments were amplified by using the primer pairs Nat1_g112-F/Nat1_g112-R, Nat1_g457-F/Nat1_g457-R, Hyg_g28F/Hyg_g28R, and Hyg_g985F/Hyg_g985R. Additional common primers, including M13_F, M13_R, P33, and P34_KU, were also used as described above.

Correct integration of the DNA cassette in transformants was confirmed by colony PCR and PCR using purified genomic DNA as the template (Fig. S1D to S1F). Using *MPK1* as an example, deletion or restoration of the ORFs was verified by using primers 4514_ORFcheckF1 and 4514_ORFcheckR1. Correct insertion of the disruption cassette at both the upstream and downstream junction was confirmed by using the primer pairs 4514_discheckF1/UnivCheck-R and UnivCheck-F/4514_discheckR1 (Fig. S1D, S1E). Similarly, correct insertion of the complement cassette was verified by using the primer pairs mpk1-compchk_F1/mpk1-compchk_R1 and Nat-compchk_F2/mpk1-compchk_R2. Because the *HYG* marker was integrated into the *AGS1-*complemented strain, Hyg-compchk_F1 was used instead of Nat-compchk_F2 (Fig. S1F).

For *AGS1,* the gene was replaced with a *NAT* marker, and in the *AGS1*-complemented strain, a complement cassette containing the target gene and *HYG* marker was inserted in place of the *NAT* marker. As a result, the *AGS1* deletion strain was nourseothricin resistant and hygromycin sensitive, whereas the complemented strain was nourseothricin sensitive and hygromycin resistant. If the *NAT* cassette had integrated at an ectopic locus during *AGS1* disruption, the complemented strain would have exhibited resistance to both nourseothricin and hygromycin. In contrast, for *MPK1*/*CHS3*/*KRE5*, the deletion strains were designed to be nourseothricin sensitive and hygromycin resistant, whereas the complemented strains were expected to be nourseothricin resistant and hygromycin sensitive. Following PCR confirmation of the expected recombination events, these genotypes were further verified by assessing the growth of the deletion and complemented strains on agar plates containing 100 µg/mL nourseothricin or 200 µg/mL hygromycin (Fig. S1G).

### 2.4. Electroporation

Electroporation was performed, with minor modifications, as described by Huang and co-workers (Huang et al., 2021). One milliliter of overnight culture suspension was added to 50 mL of YPD broth [1% (w/v) yeast extract, 2% (w/v) bacto peptone, 2% (w/v) dextrose] and shaken for 4 h at 30 °C. After rinsing of the cryptococcal cells with chilled distilled water, the cells were resuspended in 25 mL of electroporation buffer (10 mM Tris-HCl, pH 7.6, 1 mM MgCl₂, 270 mM sucrose). Dithiothreitol was added (final concentration 10 mM; if the strain could not tolerate 10 mM dithiothreitol, the concentration was reduced to 1 mM), and the suspension was incubated on ice for 20 min. The cells were then rinsed with 25 mL of electroporation buffer and resuspended in 400 µL of fresh electroporation buffer. Fifty microliters of the suspension was added to 5 µL of the DNA solution containing 1 µg Cas9 fragment, 1 µg upstream guide fragment, 1 µg downstream guide fragment, and 2 µg replacement cassette. Fifty-five microliters of the DNA–cell mixture was transferred into a chilled 2-mm-gap cuvette and then electrically pulsed with a BTX ECM600 Electro Cell Manipulator (2.5 kV / Resistance mode, 1000 V, 480Ω, 200 µF). The cells in the cuvette were harvested in 1 mL of YPD or YPD containing 1 M sorbitol and then shaken at 30 °C for 3 to 4 h. One hundred microliters each of undiluted, 10-fold diluted, and 100-fold diluted suspensions were plated onto YPD plates containing nourseothricin (JenaBioScience GmbH #96736-11-7; 100 µg/mL), G418 (Nacalai Tesque Inc.#09380-86, 200 µg/mL), or hygromycin (Nacalai Tesque Inc.#09287-84, 200 µg/mL). The plates were incubated at 30 °C for at least 2 days. Although countless colonies appeared, even at 100-fold dilution, the majority of clones were those with the correct recombination.

### 2.5. Colony PCR and PCR using purified genomic DNA

In the simplified PCR method, a suspension of disrupted colonies was used as the template. Approximately 100 mg of 0.1-mm glass beads (Yasui Kikai Corporation #YGB01) and 100 µL of TE (Tris-EDTA) buffer were added to a 2-mL screw-cap tube, and approximately four or five colonies were collected with a toothpick and suspended in the mixture. The cryptococcal cells were disrupted in a MicroSmash MS-100R cell disrupter and micro-homogenizer (Tomy Seiko Co., Ltd.) at 3,000 rpm for 5 min. For PCR, 1 µL of the bead-crushed suspension was added as the template to a 12.5- to 25-µL reaction mixture. KOD One™ PCR Master Mix -Blue-(Toyobo, KMM-201) was used for PCR, which was performed for 35 cycles.

Purified genomic DNA was also used for genotyping (Fig. S1D to S1F). For genomic DNA isolation, cryptococcal cells were cultured in 5 mL of YPD broth at 30 °C for 2 days, washed with distilled water, and collected into 2-mL screw-cap tubes. After lyophilization, approximately equal volumes of 0.1-mm glass beads were added by eye, and the cells were disrupted by using a MicroSmash MS-100R (Tomy Seiko Co., Ltd.) at 3000 rpm for 5 min. Subsequent genomic DNA extraction was performed with a *Quick*-DNA^TM^ Miniprep Kit (ZymoResearch, #D3024), with minor modifications. The powdered cells were suspended in 600 µL of genomic lysis buffer, mixed with 600 µL of chloroform, and centrifuged. The upper aqueous phase was transferred to a Zymo-Spin IICR column (ZymoResearch), and the remaining procedures were performed in accordance with the manufacturer’s instructions. Approximately 2 µg of genomic DNA was recovered from capsular strains, whereas 10 µg or more was recovered from capsule-deficient strains. For PCR, 4 ng of genomic DNA was added to a 25-µL reaction mixture containing GoTaq^®^ Green Master Mix (Promega, #M7122) (Fig. S1D to S1F).

### 2.6. Preparation of heat-killed cryptococcal cells

Some mutants that we used were unable to grow at 37 °C, making it impossible to co-culture them with dendritic cells in the immune assays. Therefore, heat-killed (HK) cryptococcal cells were used as antigenic particles in all experiments. HK cells were prepared as previously described (Ueno et al., 2021), except that the cells were cultured in YPD broth for 2 days before heat treatment.

### 2.7. Cell wall, plasma membrane, and thermal stress tests

These assays were performed as previously described (Ueno et al., 2023a). Calcofluor white (CFW), Congo red (CR), or sodium dodecyl sulfate (SDS) was added to YPD agar at the concentrations indicated in the caption to Fig. 1, and each medium, or YPD agar with no added reagent, was autoclaved (121 °C, >120 kPa, 15 to 20 min). *C. neoformans* strains were cultured in YPD broth for 2 days with shaking and then washed with PBS and adjusted to a concentration of 1 × 10^7^ cells/mL. Ten-fold serial dilutions were prepared, and 5 µL of each suspension was spotted onto each agar plate. The plates were then incubated at 30 or 37 °C for 4 days, after which colony formation was photographed with a flatbed photo and film scanner (GT-X970, Epson).

**Fig. 1.**
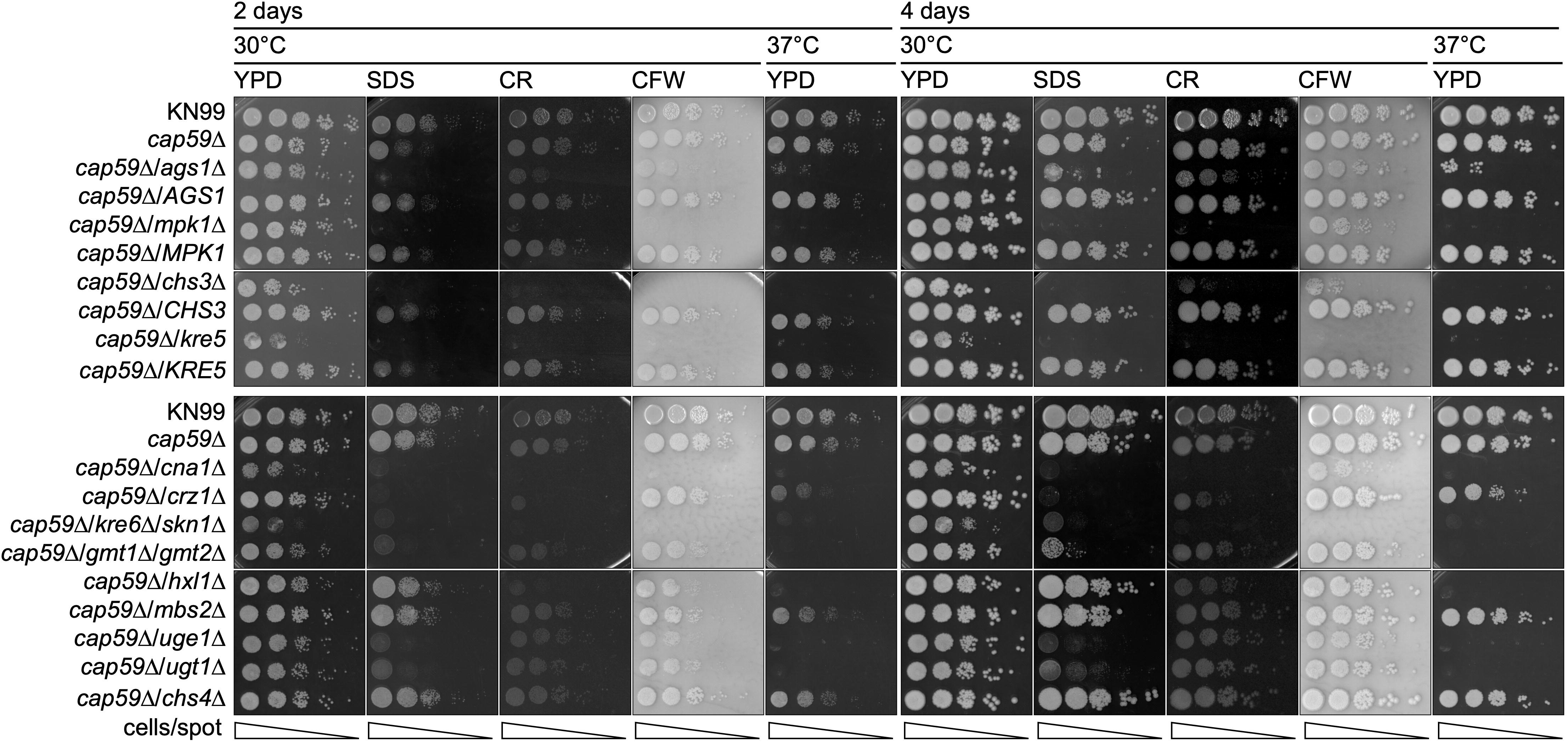
Sensitivity to cell walls, plasma membranes, and thermal stresses. *C. neoformans* strains were grown in YPD broth at 30 °C for 2 days and rinsed with PBS, then adjusted to a concentration of 1 × 10^7^ cells/mL. Tenfold serial dilutions were prepared, and 5 µL of the suspension was spotted onto YPD agar. The concentrations of each reagent were as follows: sodium dodecyl sulfate (SDS) 0.01%, Congo red (CR) 5 mg/mL, and calcofluor white (CFW) 7 mg/mL. Plates were incubated at 30 °C or 37 °C for 4 days. The experiments were conducted independently more than three times. Representative images were shown.

### 2.8. Determination of cell wall exposure

Cell wall exposure was assessed as described previously (Ueno et al., 2023a, 2019). Culture supernatant containing dectin-1-Fc, secreted by HEK293T cells transfected with expression vector, was used directly in this assay (Ueno et al., 2019). The following reagents were used: dectin-2 (Enzo Life Sciences; final concentration 2.5 µg/mL), Alexa Fluor^®^ 405 wheat germ agglutinin (WGA) (Thermo Fisher Scientific, #W56132, final concentration 10 µg/mL), anti-β-1,3-glucan monoclonal antibody (mAb) (BioSupplies Australia, #400–2, final concentration 5 µg/mL), Alexa Fluor^®^ 647 concanavalin A (ConA) (Thermo Fisher Scientific, #C21421, final concentration 50 µg/mL), Alexa Fluor^®^ 488 anti-mouse IgG polyclonal antibody (pAb) (Jackson ImmunoResearch Laboratories Inc., #115-545-071; 1:200 dilution), and Alexa Fluor^®^ 647 anti-human IgG pAb (Jackson ImmunoResearch Laboratories Inc., #709-605-149; 1:200 dilution). Hanks’s Balanced Salt Solution containing magnesium and calcium (Nacalai Tesque Inc.) was used for cell washing and staining. Flow cytometric analysis was performed with a BD FACSCanto™ II flow cytometer (BD Biosciences), and the data were analyzed with FlowJo 10 software. Fluorescent probe binding among strains was compared using the mean fluorescence intensity (MFI).

### 2.9. Measurement of 1,3BG in hot-water extraction

The supernatant was collected from HK cryptococcal suspension (1 × 10^9^ cells/mL in PBS) after incubation at 94 to 98 °C for 1 hr. The concentration of 1,3BG in the supernatant was measured with a FANGITEC^®^ G test MK-II (Shimadzu Corporation), which is based on the *Limulus* amebocyte lysate Factor G test.

### 2.10. Immune assay with bone marrow-derived dendritic cells

This assay was performed as described previously (Ueno et al., 2023a, 2021, 2019). Briefly, bone marrow-derived dendritic cells (BMDCs) were generated by differentiating bone marrow cells in the presence of 10 ng/mL granulocyte-monocyte colony-stimulating factor (GM-CSF) for 6 to 7 days. BMDCs (2 × 10^5^ cells/well) and HK cryptococci were added to flat-bottomed 96-well plates (Corning #3595) and incubated at 37°C for 24 h. After the incubation, the culture supernatants were collected, and IL-6 and IL-1β concentrations were measured by using a BioLegend ELISA MAX Standard Set (#431301 or #432601).

### 2.11. Statistics

All statistical analyses were performed with GraphPad Prism 10 software (GraphPad Software, Inc.). Differences were considered statistically significant at *P* < 0.05.

## 3. Results

### 3.1. Assessment of cell wall integrity in the series of deletion strains

From the results of previous studies, we selected target genes of which the disruption was expected to cause substantial changes in cell wall composition (Table 1). *AGS1*, *MPK1*, *CHS3*, and *KRE5* were chosen as the primary targets, and both deletion and complemented strains were generated (Fig. S1A to G). *AGS1,* which encodes 1,3AG synthase, was selected to validate the role of 1,3AG in masking 1,3BG and promoting dectin-1 evasion in *C. neoformans* (Beauvais et al., 2013; Rappleye et al., 2007; Reese et al., 2007; Reese and Doering, 2003). *MPK1* encodes a MAP (mitogen-activated protein) kinase that regulates 1,3BG exposure at 37 °C, and we investigated whether it also contributes to glucan masking under standard culture conditions at 30 °C (Bloom et al., 2021). *CHS3* encodes a homologue of chitin synthase, and deletion of this gene results in loss of chitosan, rather than chitin, from the cryptococcal cell wall (Baker et al., 2007; Banks et al., 2005). Chitosan-deficient strains are highly immunogenic and can induce excessive inflammation responses; however, whether chitosan deficiency alters 1,3BG exposure has not yet been determined (Hellmann et al., 2026; Hole et al., 2020). *KRE5* plays a pivotal role in 1,6BG synthesis in both *C. albicans* and *C. neoformans*. Although the 1,3BG content is compensatorily increased in 1,6BG-deficient strains of *C. albicans*, whether a similar compensatory response occurs in *C. neoformans* remains unknown (Gilbert et al., 2010; Herrero et al., 2004).

Disruption of these genes in the encapsulated wild-type *C. neoformans* strain KN99 revealed that the *ags1*Δ mutant was acapsular, whereas the *chs3*Δ and *kre5*Δ mutants had markedly thinner capsules than the wild type (Fig. S1H). Therefore, even if 1,3BG were exposed in these mutants and enhanced immunostimulatory activity were observed, it would be difficult to exclude the possibility that these phenotypes resulted from capsule defects rather than from changes in cell wall glucan exposure. To investigate the mechanisms underlying 1,3BG exposure independently of the capsule, and to evaluate the antigenic potential of glucan-unmasked strains, we generated a series of deletion mutants by using the acapsular *cap59*Δ as the parental background and characterized their phenotypes. In addition, deletion mutants of secondary target genes functionally associated with the primary targets were constructed and compared with the primary mutants (Tables 1 and 2).

Generally, cryptococcal strains lacking genes directly involved in cell wall integrity or cell wall synthesis are hypersensitive to cell wall, cell membrane, and thermal stress. The susceptibility of deletion strains to these stresses was examined with a spot assay (Fig. 1). The parental capsule-deficient strain *cap59*Δ lacked capsule layers (Fig. S1H), yet its susceptibility to these stresses was comparable to that of the wild-type strain KN99 (Fig. 1). In the series of deletion mutants derived from *cap59*Δ, all strains—except for *cap59*Δ/*mbs2*Δ and *cap59*Δ/*chs4*Δ—were vulnerable to either cell wall, cell membrane, or heat stress (Fig. 1). These phenotypes were consistent with the previous data (Table 1), suggesting that these genes are required for cell wall integrity.

*MBS2* is a homolog of *SWI6* in budding yeast (*Saccharomyces cerevisiae*). Swi6p in budding yeast is a transcription regulator that is phosphorylated by Mpk1p under cell wall stress conditions and regulates the transcription of *FKS2* (Kim et al., 2010; Kim and Levin, 2010). The sensitivity of *cap59*Δ/*mbs2*Δ to stress was comparable to that of *cap59*Δ (Fig. 1), suggesting that *MBS2* plays only a limited role in cell wall integrity, or there may be other compensatory mechanisms at work. (Fig. 1). Similarly, although *chs4*Δ results in a 50% reduction in chitin content (Banks et al., 2005; Rodrigues et al., 2018), *cap59*Δ/*chs4*Δ had stress tolerance comparable to *cap59*Δ (Fig. 1), suggesting that other homologous genes, namely *CHS1* to *CHS8*, may compensate for the loss of its function (Banks et al., 2005; Rodrigues et al., 2018).

### 3.2. Identification of deletion mutants with exposed cell wall antigens

In those mutant strains that showed stress hypersensitivity, their cell wall composition may have changed, possibly exposing cell wall antigens that were normally masked. Next, to assess the exposure of cell wall antigens, we used flow cytometry to analyze the extent to which the immune receptors dectin-1 and dectin-2, which recognize cell wall glucans and mannans bound to each strain (Fig. 2). Although LYSMD3 (LysM domain-containing protein 3), TLR2 (Toll-like receptor 2), TLR9 (Toll-like receptor 9), MR (mannose receptor), and NOD2 (Nucleotide-binding oligomerization domain-containing protein 2) are known to be receptors that recognize chitin, some of these receptors also cross-react with other ligands, such as glucan, mannan, and nucleic acids. However, no binding assay has been established in cryptococcal cells using their soluble receptors (Fuchs et al., 2018; He et al., 2021; Wagener et al., 2014). Because CFW binds to β-linked polysaccharides and the anionic eosin-Y interacts with a variety of cationic substances in addition to chitosan, neither is sufficiently specific for chitin or chitosan detection in this context (Monheit et al., 1984; Rieder et al., 2012; Waheed et al., 2000). Moreover, although these low-molecular-weight fluorescent dyes can be used to indirectly estimate chitin or chitosan content, they do not allow reliable assessments of chitin or chitosan surface exposure. Therefore, we used WGA, a chitin- or chitosan-binding lectin widely used for cell wall studies in cryptococcal cells, to quantify the exposure of chitin or chitosan (Fig. 2). The deposition levels of dectin-1, dectin-2, and WGA were measured via flow cytometry. In advance, the binding properties of each probe were confirmed by using *C. albicans* SC5314 as a positive control. Taking the effects of autofluorescence into account, we calculated the deposition index (DI)—the ratio of the fluorescence intensity of the fluorescent sample to that of the unstained sample—to compare the amount of receptor–protein binding for each strain (Fig. 2 and S2).

**Fig. 2.**
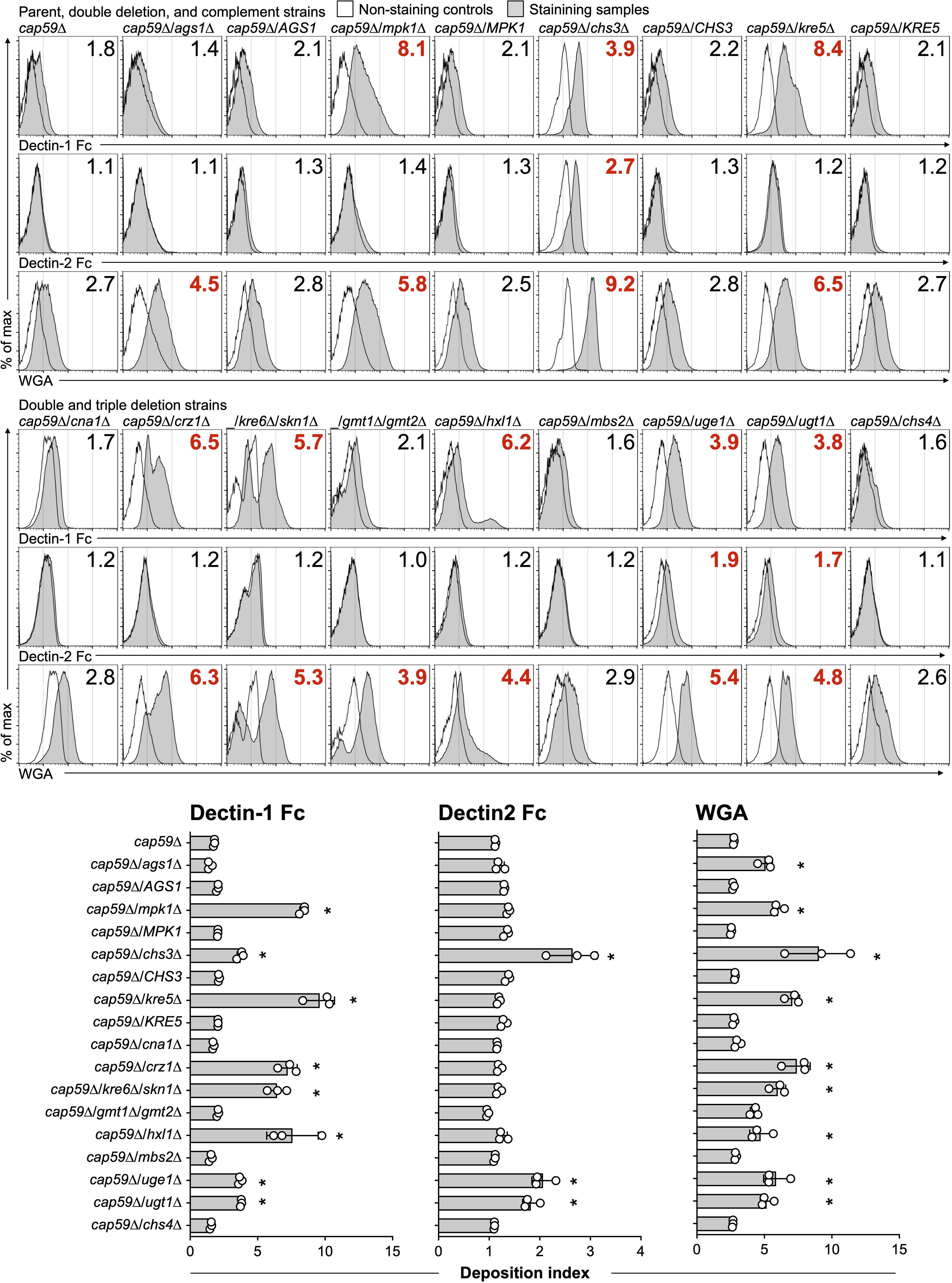
Cell wall exposure in the deletion mutants. Cryptococcal strains were cultured in YPD broth at 30 °C for 2 days and then inactivated at 94 to 98 °C for 1 hr. Deposition of dectin-1 Fc, dectin-2 Fc, and wheat germ agglutinin (WGA) was measured with flow cytometry. The values shown in the histogram represent the deposition index (DI), and red indicates a statistically significant difference from *cap59*Δ. The deposition index is defined as the ratio of the increase in the mean fluorescence intensity (MFI) when cells are labeled with each fluorescent probe, with the MFI of unlabeled cells set to one. The experiments were conducted independently three times. Representative histograms and pooled data are shown (means ± SD). \**P* < 0.05 vs. *cap59*Δ, by analysis of variance (ANOVA) with Dunnett’s *post hoc* test.

Among the primary target gene-deletion strains, *cap59*Δ/*mpk1*Δ (DI 8.1), *cap59*Δ/*chs3*Δ (DI 3.9), and *cap59*Δ/*kre5*Δ (DI 8.4) exhibited higher dectin-1 deposition than the parental strain *cap59*Δ (DI 1.8). Dectin-2 deposition was significantly increased only in *cap59*Δ/*chs3*Δ (DI 2.7 vs. 1.1 in the parent), whereas WGA binding was elevated in all primary target deletion strains (DI 4.5 to 9.2 vs. 2.7 in the parent). In contrast, the complemented strains showed deposition levels comparable to those of the parental strain *cap59*Δ. These findings do not support the previous hypothesis that 1,3BG is broadly exposed in 1,3AG-deficient strains (Beauvais et al., 2013; Rappleye et al., 2007). Although it remains unclear whether increased exposure of chitin or chitosan directly stimulates TLR2 or other pattern-recognition receptors to promote inflammation, chitin or chitosan has also been reported to induce anti-inflammatory cytokines such as IL-10 under certain conditions (Fuchs et al., 2018; He et al., 2021; Hellmann et al., 2026; Wagener et al., 2014). Nevertheless, the increased binding to dectin-1 and/or dectin-2, or both, observed in *cap59*Δ*/mpk1*Δ*, cap59*Δ/*chs3*Δ, and *cap59*Δ*/kre5*Δ suggests that these strains have reduced capacity to evade innate immune recognition via dectin receptors, potentially leading to enhanced inflammatory responses.

We next examined cell wall antigen exposure in the secondary target gene deletion strains (Fig. 2). Five strains—*cap59*Δ/*crz1*Δ (DI 6.5), *cap59*Δ/*kre6*Δ/*skn1*Δ (DI 5.7), *cap59*Δ/*hxl1*Δ (DI 6.2), *cap59*Δ/*uge1*Δ (DI 3.9), and *cap59*Δ/*ugt1*Δ (DI 3.8)—displayed higher dectin-1 deposition than the parental strain (DI 1.8). These strains also showed increased WGA binding (DI 3.9 to 6.3 vs. 2.7 in the parent). In addition, *cap59*Δ/*uge1*Δ and *cap59*Δ/*ugt1*Δ exhibited modest increases in dectin-2 deposition. Although *cap59*Δ/*gmt1*Δ/*gmt2*Δ showed no increases in dectin-1 and dectin-2 binding, WGA deposition was elevated (DI 3.9 vs. 2.8 in the parent). In contrast, *cap59*Δ/*cna1*Δ, *cap59*Δ/*mbs2*Δ, and *cap59*Δ/*chs4*Δ exhibited deposition levels of dectin-1, dectin-2, and WGA comparable to those of the parental strain (Fig. 2). These results suggest that the five deletion strains with enhanced dectin-1 binding—*cap59*Δ/*crz1*Δ, *cap59*Δ/*kre6*Δ/*skn1*Δ, *cap59*Δ/*hxl1*Δ, *cap59*Δ/*uge1*Δ, and *cap59*Δ/*ugt1*Δ—have reduced capacity to evade dectin-1-mediated innate immune recognition and may therefore elicit a relatively strong inflammatory response.

Because increased deposition of dectin-1 or dectin-2, or both, was observed in eight deletion strains, we further evaluated the binding of an anti-β-1,3-glucan mAb and the mannan-binding lectin ConA (Fig. S3). Overall, the deposition patterns of the 1,3BG mAb and ConA were consistent with those of dectin-1 and dectin-2. However, 1,3BG mAb deposition in *cap59*Δ/*uge1*Δ and *cap59*Δ/*ugt1*Δ remained comparable to that of the parental strain, despite increased dectin-1 binding (Fig. S3). This discrepancy may reflect differences in ligand recognition. Dectin-1 preferentially recognizes β-glucans containing β-1,3-linked backbones with β-1,6-linked branches of approximately 10 or more glucose residues, whereas the 1,3BG mAb recognizes shorter β-1,3-linked oligosaccharides consisting of approximately five glucose residues (Palma et al., 2006; Torosantucci et al., 2009). Furthermore, dectin-1 retains binding to cryptococcal cells following β-1,3-glucanase treatment, suggesting the presence of additional dectin-1 ligands besides 1,3BG (Ueno et al., 2019). Therefore, ligands other than 1,3BG may be exposed or enriched in *cap59*Δ/*uge1*Δ and *cap59*Δ/*ugt1*Δ, accounting for the increased dectin-1 deposition observed in these strains.

Next, in addition to evaluating 1,3BG exposure, we quantified surface-associated 1,3BG in the eight strains with increased dectin-1 deposition. To assess this, the amount of soluble 1,3BG released into the supernatant after heat treatment was quantified by using a *Limulus* amebocyte lysate Factor G assay (Fig. S4). In a pilot study, we measured the 1,3BG content of the alkaline-soluble and alkaline-insoluble cell wall fractions from cryptococcal cells. However, no notable differences were observed between these fractions. Therefore, we adopted the hot-water extraction method for 1,3BG quantification by the *Limulus amebocyte* lysate assay (Fig. S4). The amount of 1,3BG detected in the hot-water extract was generally correlated with the deposition of the anti-1,3BG mAb. The 1,3BG contents of the hot-water extracts from *cap59*Δ/*uge1*Δ and *cap59*Δ/*ugt1*Δ were comparable to those of the parent strain, *cap59*Δ. In contrast, significantly more 1,3BG was extracted from the following six deletion strains than from the parent strain: *cap59*Δ/*mpk1*Δ, *cap59*Δ/*chs3*Δ, *cap59*Δ/*kre5*Δ, *cap59*Δ/*crz1*Δ, *cap59*Δ/*kre6*Δ/*skn1*Δ, and *cap59*Δ/*hxl1*Δ (Fig. S4).

### 3.3. Determination of inflammatory potential of the deletion strains

Next, we stimulated BMDCs with the constructed deletion strains and measured IL-6 and IL-1β concentrations in the culture supernatant (Fig. 3). We plotted both representative data (Fig. 3) and pooled results data from multiple independent experiments (Fig. S5). In addition to the direct comparisons with the parent strain *cap59*Δ (Fig. 3), analysis of variance was performed to account for experimental variability and induction ratios (Fig. S5). These analyses indicated that four of the eight mutants with increased dectin-1 deposition—*cap59*Δ/*kre5*Δ, *cap59*Δ/*crz1*Δ, *cap59*Δ/*kre6*Δ/*skn1*Δ, and *cap59*Δ/*uge1*Δ—strongly induced the production of both IL-6 and IL-1β. The levels of both cytokines were significantly higher than those induced by *cap59*Δ (Figs. 3 and S5). The remaining four strains—*cap59*Δ/*mpk1*Δ, *cap59*Δ/*chs3*Δ, *cap59*Δ/*hxl1*Δ, *cap59*Δ/*uge1*Δ, and *cap59*Δ/*ugt1*Δ—stimulated BMDCs to produce significantly higher levels of either IL-6 or IL-1β than did *cap59*Δ (Figs. 3 and S5). Although *cap59*Δ/*chs4*Δ and *cap59*Δ/*mbs2*Δ induced slight increases in IL-6 or IL-1β production, these effects were much weaker than those observed for *cap59*Δ/*kre5*Δ (Figs. 3 and S5). Interestingly, *cap59*Δ/*cna1*Δ and *cap59*Δ/*gmt1*Δ/*gmt2*Δ, despite showing no obvious increases in cell wall antigen exposure, also stimulated BMDCs to produce significantly higher levels of both IL-6 and IL-1β (Figs. 3 and S5). The underlying mechanism remains unclear, although these strains may unexpectedly expose increased amounts of other pathogen-associated molecular patterns (PAMPs). Furthermore, the antigenic potential of *cap59*Δ/*ags1*Δ was not greater than that of *cap59*Δ; this findin does not support the working hypothesis that 1,3AG masks cell wall antigens to facilitate immune evasion (Beauvais et al., 2013; Rappleye et al., 2007) (Figs. 3 and S5). Overall, the eight deletion strains exhibiting increased dectin-1 deposition also showed enhanced inflammatory potential. These findings suggest that these strains are more readily recognized by dendritic cells and therefore possess reduced immune-evasion capacity.

**Fig. 3.**
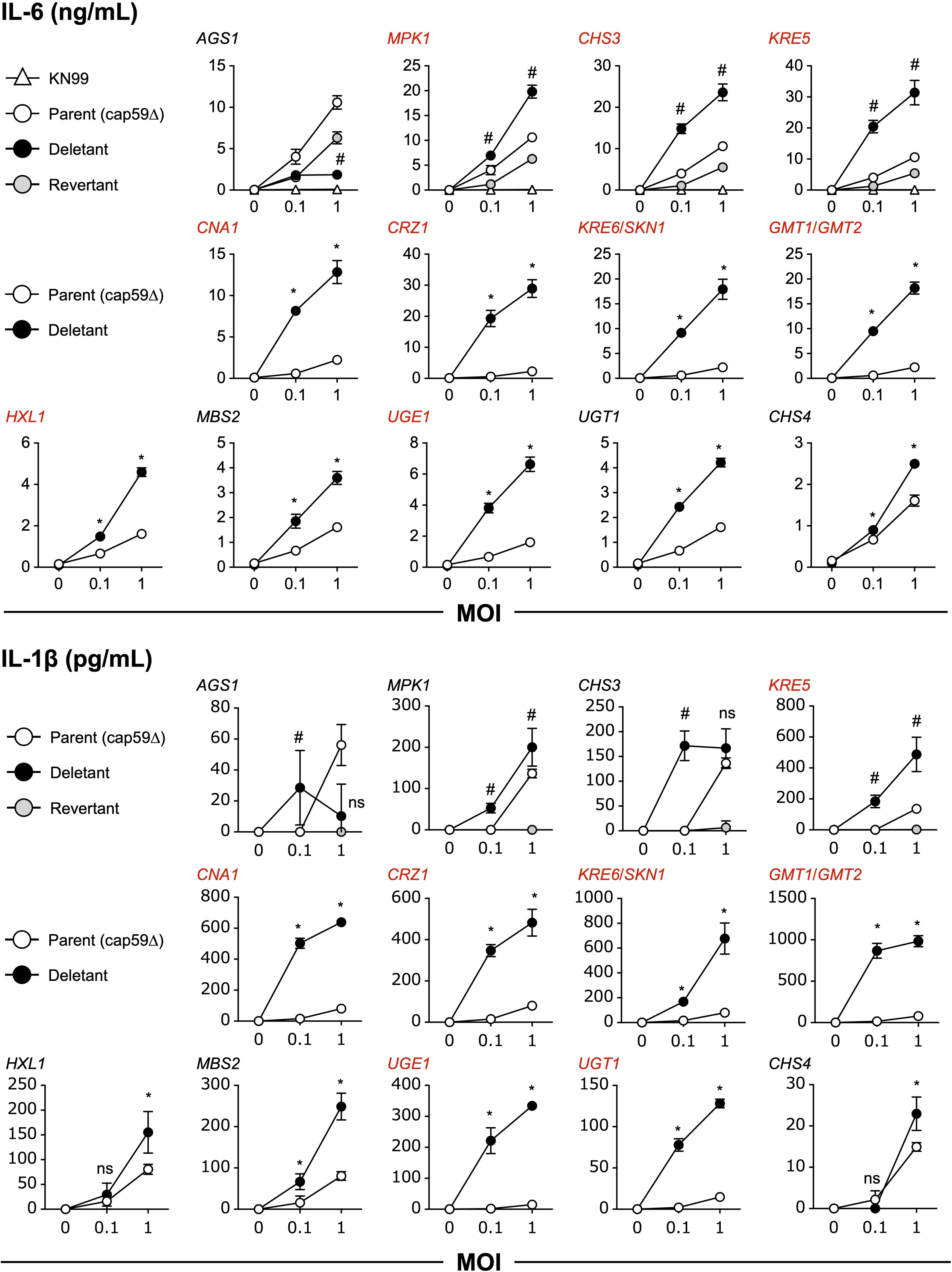
Inflammatory potential of the deletion strains. Heat-inactivated cryptococcal cells and BMDCs (2 × 10^5^ cells/well) were incubated in flat-bottomed 96-well plates for 24 h, and then cytokine levels were measured by ELISA. The experiments were conducted independently more than three times.Representative line charts are shown (means ± SD). Given the data on increased cytokine production shown in Fig. S5, strains with clearly high immunostimulatory activity are depicted in red. #*P* < 0.05 vs. *cap59*Δ and these complemented strains at the identical multiplicity of infection (MOI) point, by ANOVA with Dunnett’s *post hoc* test. \**P* < 0.05 vs. *cap59*Δ at the identical MOI, by ANOVA with Sidak’s *post hoc* test. ns: not significant.

### 3.4. Involvement of dectin-1 in enhanced inflammatory responses

To investigate the contribution of dectin-1 to the enhanced inflammatory responses induced by the deletion strains, we measured IL-6 production in BMDCs derived from dectin-1-deficient mice (Figs. 4 and S6). Compared with wild-type BMDCs, dectin-1-deficient BMDCs produced significantly smaller amounts of IL-6 when stimulated with the eight strains exhibiting increased dectin-1 deposition. In contrast, similar levels of IL-6 production were observed between wild-type and dectin-1-deficient BMDCs when stimulated with the parent strain *cap59*Δ (Fig. 4). Although the suppression levels varied among the strains, these findings indicated that dectin-1 is at least partially responsible for recognizing *C. neoformans* strains with unmasked cell wall antigens.

**Fig. 4.**
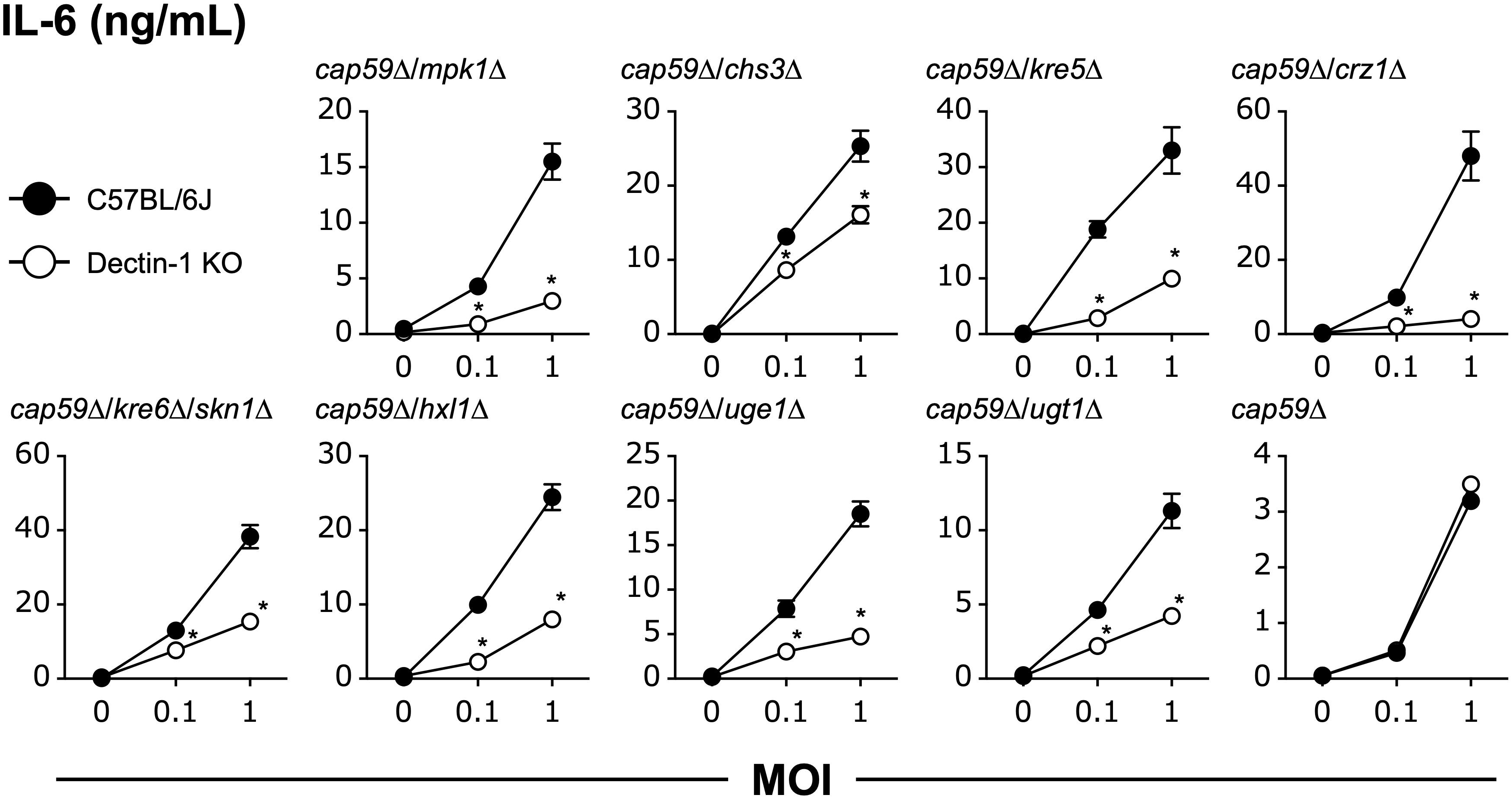
Contribution of dectin-1 to the enhanced inflammatory response induced by the deletion strains. This analysis was performed as described in the caption of Fig. 3. Representative data from two independent experiments are shown (means ± SD). \**P* < 0.05 vs. *cap59*Δ in the identical MOI, by ANOVA with Sidak’s *post hoc* test.

When dectin-1-deficient BMDCs were stimulated with *cap59*Δ/*mpk1*Δ, *cap59*Δ/*crz1*Δ, or *cap59*Δ/*ugt1*Δ, the enhanced inflammatory response was completely abolished, indicating that dectin-1 is the primary receptor responsible for recognizing these strains (Fig. S6). In contrast, *cap59*Δ/*chs3*Δ, *cap59*Δ/*kre5*Δ, *cap59*Δ/*kre6*Δ/*skn1*Δ, *cap59*Δ/*hxl1*Δ, and *cap59*Δ/*uge1*Δ still induced higher levels of IL-6 than did *cap59*Δ, even in dectin-1-deficient BMDCs (Fig. S6). These findings suggest that receptors other than dectin-1 also contribute to the recognition of these highly immunostimulatory strains (Fig. S6).

## 4. Discussion

The immune receptor dectin-1 does not bind to cryptococcal cells grown under standard laboratory conditions (YPD broth at 30 °C), even in the capsule-deficient strain *cap59*Δ, whereas dectin-1 binding is readily detected when cells are grown under host-like conditions, such as at 37 °C, in SD or Sabouraud medium (Bloom et al., 2019; Rachini et al., 2007; Ueno et al., 2019; Upadhya et al., 2023; Walsh et al., 2017). These observations indicate that cryptococcal cells strategically conceal 1,3BG under normal growth conditions to evade recognition by the dectin-1 receptor. This strategy appears to differ from those reported in other medically important fungal pathogens, including *C. albicans*, *A. fumigatus*, and *H. capsulatum* (Ballou et al., 2016; Beauvais et al., 2013; Pradhan et al., 2018; Rappleye et al., 2007). We identified the eight gene deletion strains—*cap59*Δ/*mpk1*Δ, *cap59*Δ/*chs3*Δ, *cap59*Δ/*kre5*Δ, *cap59*Δ/*crz1*Δ, *cap59*Δ/*kre6*Δ/*skn1*Δ, *cap59*Δ/*hxl1*Δ, *cap59*Δ/*uge1*Δ, and *cap59*Δ/*ugt1*Δ—that failed to evade dectin-1 recognition and consequently elicited enhanced cytokine production by innate immune cells. These strains represent promising whole-cell antigens for vaccine development against cryptococcosis, as similar strategies have proven successful in other fungal vaccine studies (Ueno et al., 2023b). A graphic summary of our findings is presented in Figure S7.

Our results indicate that the Mpk1/Crz1 pathway, the unfolded protein response, and the biosynthesis of chitosan/1,6BG/GXMGal (glucuronoxylomannogalactan) are required for capsule-independent regulation of 1,3BG exposure and dectin-1 evasion in *C. neoformans* under standard culture conditions. Because disruption of these pathways increased susceptibility to cell wall stress, these metabolic processes were essential for maintaining cell wall integrity. Our findings further suggest that intact cell wall architecture is required to conceal 1,3BG from immune recognition. Surprisingly, although 1,3AG/mannan synthesis and the calcineurin pathway also contributed to cell wall integrity, defects in these pathways did not necessarily increase dectin-1 deposition. Thus, impaired cell wall integrity alone is insufficient to explain enhanced 1,3BG exposure. The molecular mechanism underlying increased dectin-1 recognition remains unclear. One possibility is that compensatory remodeling increases the amount of surface-exposed 1,3BG in response to defects in cell wall synthesis. Alternatively, alterations in cell wall architecture may change the spatial organization or accessibility of existing 1,3BG without increasing its overall abundance. Future studies should determine whether enhanced dectin-1 binding results primarily from increased surface exposure or increased synthesis of, or structural alterations in, 1,3BG, such as changes in polymer length, branching, or triple helical organization (Anaya et al., 2023; Manabe and Yamaguchi, 2021).

On the basis of studies of *A. fumigatus* and *H. capsulatum*, we initially hypothesized that 1,3AG masks 1,3BG, thereby promoting immune evasion in *C. neoformans* (Beauvais et al., 2013; Rappleye et al., 2007). However, our findings did not support this hypothesis. Given that 1,3AG constitutes approximately 70% of the cryptococcal cell wall (Ankur et al., 2025), and in that *cap59*Δ/*ags1*Δ exhibited lower inflammatory potential than *cap59*Δ, it is possible that loss of 1,3AG disrupted the structural scaffold of the cell wall, resulting in leakage or loss of cell wall antigens, rather than enhanced antigen exposure. Consistent with this interpretation, the amount of alkali-soluble 1,3BG in *cap59*Δ/*ags1*Δ is less than half that detected in *cap59*Δ (Reese et al., 2007).

Although dectin-1/2 deposition on *cap59*Δ/*cna1*Δ and *cap59*Δ/*gmt1*Δ/*gmt2*Δ was comparable to that observed for *cap59*Δ, both strains induced significantly stronger inflammatory responses in BMDCs. Because these mutants were also hypersensitive to cell wall stress, they may expose PAMPs other than dectin-1/2 ligands. The identity of these ligands and their corresponding host receptors remains unknown. We previously demonstrated that the integrin CD11b directly recognizes the capsule-deficient strain *cap59*Δ grown aerobically in YPD broth at 30 °C, whereas GXM-coated cells are no longer recognized. Furthermore, blocking of CD11b with the monoclonal antibody M1/70 abolishes the recognition of *cap59*Δ by BMDCs (Ueno et al., 2021) (Fig. S7). Because several deletion strains retained enhanced inflammatory activity even in dectin-1-deficient BMDCs, additional receptors such as CD11b are likely to contribute to their recognition. Further studies are needed to clarify the relative contributions of these receptors, including CD11b.

Further *in vivo* studies will be required to determine whether the deletion strains identified here can serve as vaccine strains. *chs3*Δ elicits excessive pulmonary inflammation in mice, and even heat-killed cells induce lethal immunopathology (Hole et al., 2020). Therefore, careful optimization of the vaccine dose and route of administration will be essential for developing vaccines based on highly immunostimulatory strains (Hellmann et al., 2026). We previously demonstrated that *cap59*Δ^SD^, cultured in SD medium and exhibiting increased 1,3BG exposure, functions as an effective strain in vaccine development (Ueno et al., 2025). The highly immunostimulatory strains identified here induced IL-6 production at levels comparable to those elicited by *cap59*Δ^SD^, suggesting that they possess sufficient adjuvant properties. However, it remains unknown whether these strains express adequate levels of protective T cell antigens. Lung resident memory T cells induced by *cap59*Δ^SD^ recognized the chitin deacetylase Cda2, indicating that this vaccine strain expresses sufficient amounts of this protective antigen (Ueno et al., 2025). In addition to Cda2, several other protective antigens, including Blp4 (barwin-like domain protein 4), have been identified (Hester et al., 2020). Further studies should determine whether these antigens are similarly expressed in the highly immunostimulatory deletion strains identified here.

In summary, we identified eight deletion strains—*cap59*Δ/*mpk1*Δ, *cap59*Δ/*chs3*Δ, *cap59*Δ/*kre5*Δ, *cap59*Δ/*crz1*Δ, *cap59*Δ/*kre6*Δ/*skn1*Δ, *cap59*Δ/*hxl1*Δ, *cap59*Δ/*uge1*Δ, and *cap59*Δ/*ugt1*Δ—that exhibited increased dectin-1 deposition under standard growth conditions (YPD 30 °C under aerobic conditions). These strains displayed enhanced immunostimulatory activity and induced robust production of IL-6 or IL-1β, or both, by BMDCs, at least in part through dectin-1 mediated recognition. Collectively, these findings provide new insights into the capsule-independent mechanisms that regulate 1,3BG exposure and immune evasion in *C. neoformans* and identify promising candidates for the development of whole-cell vaccines against cryptococcosis.

## Declaration of competing interest

The authors declare that they have no known competing financial interests or personal relationships that could have influenced the work reported in this paper.

## Acknowledgements

This study was supported by the Japan Society for the Promotion of Science (KAKENHI 26K100125 and 23K27411), the Japan Agency for Medical Research and Development (AMED 26fk0108758), the Takeda Science Foundation, Ichiro Kanehara Foundation, Morinomiyako Medical Research Foundation, and the Shionogi Infectious Disease Research Promotion Foundation. K. Kwon-Chung was supported by intramural research funds from the National Institute of Allergy and Infectious Diseases. The authors acknowledge Dr. Shinobu Saijo (Chiba University, Japan) for kindly sharing the dectin-1 knockout mice.

## CRediT authorship contribution statement

Conceptualization: K.U.; Methodology: K.U.; Investigation: K.U., A.N., N.H., D.Y., K.M., A.K.; Formal analysis: K.U., A.N., N.H., D.Y., K.M., A.K.; Writing – original draft: K.U., K.K.C.; Writing – review & editing: K.U., K.K.C., Y.M.; Supervision: K.U., K.K.C., Y.M.

## Supplementary figure legends

**Fig. S1. Construction of cryptococcal gene-knockout strains by using a CRISPR-Cas9 system. (A) Linear DNA fragments for introduction into cryptococcal cells by using electroporation.** We adopted a strategy of using two guide sequences for a single target gene to cleave at two sites. The following four fragments were used for electroporation: the Cas9 fragment, upstream guide fragment, downstream fragment, and replacement cassette. Half-arrows indicate the relative annealing sites of the primers described in the “Materials and Methods” section. The guide fragment was amplified by two rounds of overlap extension PCR. Blue and black arrows represent the primers used in the first and second rounds, respectively. Gene-specific guide sequences are indicated by red bars and were obtained from http://grna.ctegd.uga.edu. The replacement cassettes contain the neomycin (Neo), nourseothricin (Nat), or hygromycin (Hyg) resistance genes, which were amplified from the plasmids. Green bars represent the 56-bp gene-specific homologous region adjacent to the site of cleavage by Cas9. For the complementation, *YFG* (your favorite gene) has been cloned into a plasmid in advance, and its nucleotide sequence has been verified by sequencing. The complement cassette was amplified from a plasmid carrying *YFG*. The orange arrow is a gene-specific primer that differs for each target gene. The downstream primer is universal primer 1562-compR3, which anneals to the *ACT1* promoter. **(B) Disruption.** Red arrowheads indicate the Cas9 cleavage site, and green bars are 56-bp homologous tags for the recombination. Correct integration of the DNA cassette in transformants was confirmed by PCR using the three primer sets represented by the blue arrows. **(C) Complementation.** Homologous recombination occurs at the promoter region of *YFG* and the drug-resistance gene. Their lengths are 1 kb and 0.8 kb, respectively. Correct integration of the complement cassette was confirmed by PCR using the three primer sets indicated by the blue arrows. **(D–F) Genotyping.** Deletion and restoration of ORFs, and the correct integration of the DNA cassette at both the upstream (left) and downstream (right) regions, were confirmed by PCR using three primer sets. Although colony PCR was used for routine screening, purified genomic DNA was used for these tests. M: 1-kb DNA ladder, P: parent strain KN99 or *cap59*Δ, R: revertant strain. **(G) Replacement of drug-resistance genes.** As the drug-resistance genes in the deletion mutants were replaced by the complement cassette, nourseothricin resistance was abolished in the *AGS1*-complemented strain, and hygromycin resistance was abolished in the *MPK1*/*CHS3*/*KRE5*-complemented strain. **(H) Capsule formation in the single deletion strains.** Cryptococcal cells were cultured in YPD broth at 30 °C for 2 days. Fifty microliters of the suspension and 5 mL of RPMI medium without supplements were added to a 25-cm^2^ cell-culture flask and incubated for an additional 2 days at 30 °C. One milliliter of the suspension was harvested, resuspended in Indian ink, and observed under a light microscope (Olympus IX81). Representative pictures from three independent experiments are shown.

**Fig. S2. Titer check for dectin-1 Fc, dectin-2 Fc, and WGA.** The binding level of each fluorescent probe was measured as shown in Fig. 2. Dectin-2 Fc deposition is fundamentally weak, even in the positive control *C. albicans* SC5314, and the histogram right shift is marginal. Both dectin-1 and dectin-2 Fc generally do not bind to *Cryptococcus* spp., irrespective of capsule formation. Representative histograms and a pooled graph from three independent experiments are shown (means ± SD).

**Fig. S3. Binding of anti-1,3BG mAb and ConA to the eight deletion strains with increased dectin-1 deposition.** The experiment was performed as described in the caption to Fig. 2. Representative histograms and pooled data from three independent experiments are shown (means ± SD). \**P* < 0.05 vs. *cap59*Δ, by ANOVA with Dunnett’s *post hoc* test.

**Fig. S4. Measurement of 1,3BG contents in the hot-water extract.** Cryptococcal cells were cultured in YPD broth for 2 days, then rinsed and resuspended in PBS (1 × 10^9^ cells/mL). After 1 h of boiling, the supernatant was collected to measure the 1,3BG content by using the *Limulus* amebocyte lysate Factor G test. Pooled data from four independent experiments are shown (means ± SD). \**P* < 0.05 vs. *cap59*Δ, by ANOVA with Dunnett’s *post hoc* test. #*P* < 0.05 vs. *cap59*Δ by Mann-Whitney test, but not a significantly different by ANOVA with Dunnett’s *post hoc* test. ns: not significant in any statistical analysis.

**Fig. S5. Induction ratios of IL-6 (A) and IL-1**β **production (B).** Left panels show pooled raw data from more than three experiments, and right panels show the induction ratios of IL-6 (A) and IL-1β (B) relative to the cytokine level upon stimulation with *cap59*Δ, set to 1. \**P* < 0.05 vs. *cap59*Δ, by ANOVA with Dunnett’s *post hoc* test. †: Ratio was not calculated because IL-1β production was not observed in some cases.

**Fig. S6. Several deletion strains still increased levels of IL-6, even in dectin-1-deficient BMDCs.** This experiment was the same as that in Fig. 3. Pooled bar graphs are shown from two independent experiments (means ± SD). \**P* < 0.05 vs. *cap59*Δ, by ANOVA with Dunnett’s *post hoc* test.

**Fig. S7. Genes for capsule-independent 1,3BG masking in *C. neoformans*.** Under thermal stress, *CCR4* plays a role in mRNA degradation, and the deficiency leads to transcriptional reprogramming failure and exposes 1,3BG (Bloom et al., 2019). In contrast, we identified novel genes that regulate 1,3BG exposure under standard culture conditions (YPD, 30 °C, aerobic). The wild-type strain KN99 is not recognized by immune cells, such as dendritic cells, without opsonization. Capsule-deficient strains are recognized by the integrin CD11b on dendritic cells but not by dectin-1 (Ueno et al., 2021, 2019; Upadhya et al., 2023). Deletion strains with increased dectin-1 deposition stimulated BMDCs to produce significantly higher levels of IL-6 and IL-1β than in the parent strain. Although this hyperinflammation was mediated by dectin-1 in BMDCs, several deletion strains still induced higher amounts of IL-6 even in dectin-1-deficient BMDCs. Therefore, it is possible that receptors other than dectin-1 also contribute to hyperinflammation.

## Notes

### Competing Interest Statement

The authors have declared no competing interest.

### Summary of Updates

This version has been proofread and edited by an English editing service.

## References

Anaya, E.U., Amin, A.E., Wester, M.J., Danielson, M.E., Michel, K.S., Neumann, A.K., 2023. Dectin-1 multimerization and signaling depends on fungal β-glucan structure and exposure. Biophys. J. 122, 3749–3767. 10.1016/j.bpj.2023.07.021

Ankur, A., Yarava, J.R., Gautam, I., Scott, F.J., Mentink-Vigier, F., Chrissian, C., Xie, L., Roy, D., Stark, R.E., Doering, T.L., Wang, P., Wang, T., 2025. Polymorphic α-Glucans as structural scaffolds in *Cryptococcus* cell walls for chitin, capsule, and melanin: Insights from ^13^C and ^1^H solid-state NMR. Angew. Chem. Int. Ed. e202510409. 10.1002/anie.202510409

Baker, L.G., Specht, C.A., Donlin, M.J., Lodge, J.K., 2007. Chitosan, the deacetylated form of chitin, is necessary for cell wall integrity in *Cryptococcus neoformans*. Eukaryot Cell 6, 855–867. 10.1128/ec.00399-06

Ballou, E.R., Avelar, G.M., Childers, D.S., Mackie, J., Bain, J.M., Wagener, J., Kastora, S.L., Panea, M.D., Hardison, S.E., Walker, L.A., Erwig, L.P., Munro, C.A., Gow, N.A.R., Brown, G.D., MacCallum, D.M., Brown, A.J.P., 2016. Lactate signalling regulates fungal β-glucan masking and immune evasion. Nat Microbiol 2, 16238. 10.1038/nmicrobiol.2016.238

Banks, I.R., Specht, C.A., Donlin, M.J., Gerik, K.J., Levitz, S.M., Lodge, J.K., 2005. A chitin synthase and its regulator protein are critical for chitosan production and growth of the fungal pathogen *Cryptococcus neoformans*. Eukaryot. Cell 4, 1902–1912. 10.1128/ec.4.11.1902-1912.2005

Beauvais, A., Bozza, S., Kniemeyer, O., Formosa, Cécile, Formosa, Céline, Balloy, V., Henry, C., Roberson, R.W., Dague, E., Chignard, M., Brakhage, A.A., Romani, L., Latgé, J.-P., 2013. Deletion of the α-(1,3)-glucan synthase genes induces a restructuring of the conidial cell wall responsible for the avirulence of *Aspergillus fumigatus*. PLoS Pathog. 9, e1003716. 10.1371/journal.ppat.1003716

Bloom, A.L.M., Goich, D., Knowles, C.M., Panepinto, J.C., 2021. Glucan unmasking identifies regulators of temperature-induced translatome reprogramming in *C. neoformans*. mSphere 6. 10.1128/msphere.01281-20

Bloom, A.L.M., Jin, R.M., Leipheimer, J., Bard, J.E., Yergeau, D., Wohlfert, E.A., Panepinto, J.C., 2019. Thermotolerance in the pathogen *Cryptococcus neoformans* is linked to antigen masking via mRNA decay-dependent reprogramming. Nat Commun 10, 4950–13. 10.1038/s41467-019-12907-x

Chang, Y.C., Kwon-Chung, K.J., 1994. Complementation of a capsule-deficient mutation of *Cryptococcus neoformans* restores its virulence. Mol. Cell. Biol. 14, 4912–4919. 10.1128/mcb.14.7.4912-4919.1994

Cheon, S.A., Jung, K.-W., Chen, Y.-L., Heitman, J., Bahn, Y.-S., Kang, H.A., 2011. Unique evolution of the UPR pathway with a novel bZIP transcription factor, Hxl1, for controlling pathogenicity of *Cryptococcus neoformans*. PLoS Pathog 7, e1002177. 10.1371/journal.ppat.1002177

Denham, S., Brown, J., 2018. Mechanisms of pulmonary escape and dissemination by *Cryptococcus neoformans*. JoF 4, 25–17. 10.3390/jof4010025

Diniz-Lima, I., Fonseca, L.M. da, Silva-Junior, E.B. da, Guimarães-de-Oliveira, J.C., Freire-de-Lima, L., Nascimento, D.O., Morrot, A., Previato, J.O., Mendonça-Previato, L., Decote-Ricardo, D., Freire-de-Lima, C.G., 2022. *Cryptococcus*: history, epidemiology and immune evasion. Appl. Sci. 12, 7086. 10.3390/app12147086

Esher, S.K., Ost, K.S., Kohlbrenner, M.A., Pianalto, K.M., Telzrow, C.L., Campuzano, A., Nichols, C.B., Munro, C., Wormley, F.L., Alspaugh, J.A., 2018. Defects in intracellular trafficking of fungal cell wall synthases lead to aberrant host immune recognition. PLoS Pathog 14, e1007126. 10.1371/journal.ppat.1007126

Farkas, V., Takeo, K., Maceková, D., Ohkusu, M., Yoshida, S., Sipiczki, M., 2009. Secondary cell wall formation in *Cryptococcus neoformans* as a rescue mechanism against acid-induced autolysis. FEMS yeast research 9, 311–320. 10.1111/j.1567-1364.2008.00478.x

Fraser, J.A., Subaran, R.L., Nichols, C.B., Heitman, J., 2003. Recapitulation of the sexual cycle of the primary fungal pathogen *Cryptococcus neoformans* var. *gattii*: implications for an outbreak on Vancouver Island, Canada. Eukaryot Cell 2, 1036–1045.

Fuchs, K., Gloria, Y.C., Wolz, O.-O., Herster, F., Sharma, L., Dillen, C.A., Täumer, C., Dickhöfer, S., Bittner, Z., Dang, T.-M., Singh, A., Haischer, D., Schlöffel, M.A., Koymans, K.J., Sanmuganantham, T., Krach, M., Roger, T., Roy, D.L., Schilling, N.A., Frauhammer, F., Miller, L.S., Nürnberger, T., LeibundGut-Landmann, S., Gust, A.A., Macek, B., Frank, M., Gouttefangeas, C., Cruz, C.S.D., Hartl, D., Weber, A.N., 2018. The fungal ligand chitin directly binds TLR2 and triggers inflammation dependent on oligomer size. EMBO Rep 19, e46065. 10.15252/embr.201846065

Garcia-Rubio, R., Oliveira, H.C. de, Rivera, J., Trevijano-Contador, N., 2020. The fungal cell wall: *Candida*, *Cryptococcus*, and *Aspergillus* species. Front. Microbiol. 10, 2993. 10.3389/fmicb.2019.02993

Gilbert, N.M., Donlin, M.J., Gerik, K.J., Specht, C.A., Djordjevic, J.T., Wilson, C.F., Sorrell, T.C., Lodge, J.K., 2010. *KRE* genes are required for β-1,6-glucan synthesis, maintenance of capsule architecture and cell wall protein anchoring in *Cryptococcus neoformans*. Mol. Microbiol. 76, 517–534. 10.1111/j.1365-2958.2010.07119.x

He, X., Howard, B.A., Liu, Y., Neumann, A.K., Li, L., Menon, N., Roach, T., Kale, S.D., Samuels, D.C., Li, H., Kite, T., Kita, H., Hu, T.Y., Luo, M., Jones, C.N., Okaa, U.J., Squillace, D.L., Klein, B.S., Lawrence, C.B., 2021. LYSMD3: A mammalian pattern recognition receptor for chitin. Cell Rep 36, 109392. 10.1016/j.celrep.2021.109392

Hellmann, M.J., Upadhya, R., Tchoub, E., Moerschbacher, B.M., Lodge, J.K., Cord-Landwehr, S., 2026. Acetylation and accessibility of *Cryptococcus neoformans* cell wall chitosans influence the strength of host immune responses. Cell Surf. 15, 100175. 10.1016/j.tcsw.2026.100175

Herrero, A.B., Magnelli, P., Mansour, M.K., Levitz, S.M., Bussey, H., Abeijon, C., 2004. *KRE5* gene null mutant strains of *Candida albicans* are avirulent and have altered cell wall composition and hypha formation properties. Eukaryot. Cell 3, 1423–1432. 10.1128/ec.3.6.1423-1432.2004

Hester, M.M., Lee, C.K., Abraham, A., Khoshkenar, P., Ostroff, G.R., Levitz, S.M., Specht, C.A., 2020. Protection of mice against experimental cryptococcosis using glucan particle-based vaccines containing novel recombinant antigens. Vaccine 38, 620–626. 10.1016/j.vaccine.2019.10.051

Hole, C.R., Lam, W.C., Upadhya, R., Lodge, J.K., 2020. *Cryptococcus neoformans* chitin synthase 3 plays a critical role in dampening host inflammatory responses. mBio 11. 10.1128/mbio.03373-19

Huang, M.Y., Joshi, M.B., Boucher, M.J., Lee, S., Loza, L.C., Gaylord, E.A., Doering, T.L., Madhani, H.D., 2021. Short homology-directed repair using optimized Cas9 in the pathogen *Cryptococcus neoformans* enables rapid gene deletion and tagging. Genetics 220, iyab180. 10.1093/genetics/iyab180

James, P.G., Cherniak, R., Jones, R.G., Stortz, C.A., Reiss, E., 1990. Cell-wall glucans of *Cryptococcus neoformans* CAP 67. Carbohydr. Res. 198, 23–38. 10.1016/0008-6215(90)84273-w

Jung, K.-W., Yang, D.-H., Maeng, S., Lee, K.-T., So, Y.-S., Hong, J., Choi, J., Byun, H.-J., Kim, H., Bang, S., Song, M.-H., Lee, J.-W., Kim, M.S., Kim, S.-Y., Ji, J.-H., Park, G., Kwon, H., Cha, S., Meyers, G.L., Wang, L.L., Jang, J., Janbon, G., Adedoyin, G., Kim, T., Averette, A.K., Heitman, J., Cheong, E., Lee, Y.-H., Lee, Y.-W., Bahn, Y.-S., 2015. Systematic functional profiling of transcription factor networks in *Cryptococcus neoformans*. Nat Commun 6, 6757–14. 10.1038/ncomms7757

Kim, K.-Y., Levin, D.E., 2010. Transcriptional reporters for genes activated by cell wall stress through a non-catalytic mechanism involving Mpk1 and SBF. Yeast (Chichester, Engl.) 27, 541–8. 10.1002/yea.1782

Kim, K.-Y., Truman, A.W., Caesar, S., Schlenstedt, G., Levin, D.E., 2010. Yeast Mpk1 cell wall integrity mitogen-activated protein kinase regulates nucleocytoplasmic shuttling of the Swi6 transcriptional regulator. Mol. Biol. Cell 21, 1609–1619. 10.1091/mbc.e09-11-0923

Lev, S., Desmarini, D., Chayakulkeeree, M., Sorrell, T.C., Djordjevic, J.T., 2012. The Crz1/Sp1 transcription factor of *Cryptococcus neoformans* is activated by calcineurin and regulates cell wall integrity. PLoS ONE 7, e51403. 10.1371/journal.pone.0051403

Li, Y., Chadwick, B., Pham, T., Xie, X., Lin, X., 2024. Aspartyl peptidase May1 induces host inflammatory response by altering cell wall composition in the fungal pathogen *Cryptococcus neoformans*. mBio 15, e00920–24. 10.1128/mbio.00920-24

Manabe, N., Yamaguchi, Y., 2021. 3D structural insights into β-glucans and their binding proteins. Int. J. Mol. Sci. 22, 1578. 10.3390/ijms22041578

Monheit, J.E., Cowan, D.F., Moore, D.G., 1984. Rapid detection of fungi in tissues using calcofluor white and fluorescence microscopy. Arch. Pathol. Lab. Med. 108, 616–8.

Moyrand, F., Fontaine, T., Janbon, G., 2007. Systematic capsule gene disruption reveals the central role of galactose metabolism on *Cryptococcus neoformans* virulence. Mol. Microbiol. 64, 771–781. 10.1111/j.1365-2958.2007.05695.x

Nielsen, K., Cox, G.M., Wang, P., Toffaletti, D.L., Perfect, J.R., Heitman, J., 2003. Sexual cycle of *Cryptococcus neoformans* var. *grubii* and virulence of congenic *a* and ⍰ isolates. Infect Immun 71, 4831–4841. 10.1128/iai.71.9.4831-4841.2003

Odom, A., Muir, S., Lim, E., Toffaletti, D.L., Perfect, J., Heitman, J., 1997. Calcineurin is required for virulence of *Cryptococcus neoformans*. EMBO J. 16, 2576–2589. 10.1093/emboj/16.10.2576

O’Meara, T.R., Holmer, S.M., Selvig, K., Dietrich, F., Alspaugh, J.A., 2013. *Cryptococcus neoformans* Rim101 is associated with cell wall remodeling and evasion of the host immune responses. mBio 4, 525. 10.1128/mbio.00522-12

Palma, A.S., Feizi, T., Zhang, Y., Stoll, M.S., Lawson, A.M., Díaz-Rodríguez, E., Campanero-Rhodes, M.A., Costa, J., Gordon, S., Brown, G.D., Chai, W., 2006. Ligands for the β-glucan receptor, dectin-1, assigned using “designer” microarrays of oligosaccharide probes (neoglycolipids) generated from glucan polysaccharides*. J. Biol. Chem. 281, 5771–5779. 10.1074/jbc.m511461200

Pradhan, A., Avelar, G.M., Bain, J.M., Childers, D.S., Larcombe, D.E., Netea, M.G., Shekhova, E., Munro, C.A., Brown, G.D., Erwig, L.P., Gow, N.A.R., Brown, A.J.P., 2018. Hypoxia promotes immune evasion by triggering β-glucan masking on the *Candida albicans* cell surface via mitochondrial and cAMP-protein kinase A signaling. mBio 9, 165rv13. 10.1128/mbio.01318-18

Rachini, A., Pietrella, D., Lupo, P., Torosantucci, A., Chiani, P., Bromuro, C., Proietti, C., Bistoni, F., Cassone, A., Vecchiarelli, A., 2007. An anti-beta-glucan monoclonal antibody inhibits growth and capsule formation of *Cryptococcus neoformans in vitro* and exerts therapeutic, anticryptococcal activity *in vivo*. Infect Immun 75, 5085–5094. 10.1128/iai.00278-07

Rappleye, C.A., Eissenberg, L.G., Goldman, W.E., 2007. *Histoplasma capsulatum* α-(1,3)-glucan blocks innate immune recognition by the β-glucan receptor. Proc. Natl. Acad. Sci. 104, 1366–1370. 10.1073/pnas.0609848104

Reese, A.J., Doering, T.L., 2003. Cell wall alpha-1,3-glucan is required to anchor the *Cryptococcus neoformans* capsule. Mol. Microbiol. 50, 1401–1409.

Reese, A.J., Yoneda, A., Breger, J.A., Beauvais, A., Liu, H., Griffith, C.L., Bose, I., Kim, M.-J., Skau, C., Yang, S., Sefko, J.A., Osumi, M., Latge, J.-P., Mylonakis, E., Doering, T.L., 2007. Loss of cell wall alpha(1-3) glucan affects *Cryptococcus neoformans* from ultrastructure to virulence. Mol. Microbiol. 63, 1385–1398. 10.1111/j.1365-2958.2006.05551.x

Rieder, A., Knutsen, S.H., Ballance, S., Grimmer, S., Airado-Rodríguez, D., 2012. Cereal β-glucan quantification with calcofluor-application to cell culture supernatants. Carbohydr. Polym. 90, 1564–1572. 10.1016/j.carbpol.2012.07.031

Rodrigues, J., Ramos, C.L., Frases, S., Godinho, R.M. da C., Fonseca, F.L., Rodrigues, M.L., 2018. Lack of chitin synthase genes impacts capsular architecture and cellular physiology in *Cryptococcus neoformans*. Cell Surf 2, 14–23. 10.1016/j.tcsw.2018.05.002

Saijo, S., Fujikado, N., Furuta, T., Chung, S., Kotaki, H., Seki, K., Sudo, K., Akira, S., Adachi, Y., Ohno, N., Kinjo, T., Nakamura, K., Kawakami, K., Iwakura, Y., 2007. Dectin-1 is required for host defense against *Pneumocystis carinii* but not against *Candida albicans*. Nat. Immunol. 8, 39–46. 10.1038/ni1425

Torosantucci, A., Chiani, P., Bromuro, C., Bernardis, F.D., Palma, A.S., Liu, Y., Mignogna, G., Maras, B., Colone, M., Stringaro, A., Zamboni, S., Feizi, T., Cassone, A., 2009. Protection by ânti-β-glucan antibodies is associated with restricted β-1,3-glucan binding specificity and inhibition of fungal growth and adherence. PLoS ONE 4, e5392. 10.1371/journal.pone.0005392

Ueno, K., Nagamori, A., Honkyu, N.O., Kataoka, M., Shimizu, K., Chang, Y.C., Kwon-Chung, K.J., Miyazaki, Y., 2023a. *Cryptococcus neoformans* requires the *TVF1* gene for thermotolerance and virulence. Med. Mycol. 61, myad101. 10.1093/mmy/myad101

Ueno, K., Nagamori, A., Honkyu, N.O., Kwon-Chung, K.J., Miyazaki, Y., 2025. Lung-resident memory Th2 cells regulate pulmonary cryptococcosis by inducing type-II granuloma formation. Mucosal Immunol. 10.1016/j.mucimm.2025.02.004

Ueno, K., Otani, Y., Yanagihara, N., Nakamura, T., Shimizu, K., Yamagoe, S., Miyazaki, Y., 2019. *Cryptococcus gattii* alters immunostimulatory potential in response to the environment. PLoS ONE 14, e0220989. 10.1371/journal.pone.0220989

Ueno, K., Otani, Y., Yanagihara, N., Urai, M., Nagamori, A., Sato-Fukushima, M., Shimizu, K., Saito, N., Miyazaki, Y., 2021. *Cryptococcus gattii* evades CD11b-mediated fungal recognition by coating itself with capsular polysaccharides. Eur J Immunol. 10.1002/eji.202049042

Ueno, K., Tsuge, S., Shimizu, K., Miyazaki, Y., 2023b. Promising whole-cell vaccines against cryptococcosis. Microbiol. Immunol. 67, 211–223. 10.1111/1348-0421.13056

Upadhya, R., Lam, W.C., Hole, C.R., Vasselli, J.G., Lodge, J.K., 2023. Cell wall composition in *Cryptococcus neoformans* is media dependent and alters host response, inducing protective immunity. Front. Fungal Biol. 4, 1183291. 10.3389/ffunb.2023.1183291

Wagener, J., Malireddi, R.K.S., Lenardon, M.D., Köberle, M., Vautier, S., MacCallum, D.M., Biedermann, T., Schaller, M., Netea, M.G., Kanneganti, T.-D., Brown, G.D., Brown, A.J.P., Gow, N.A.R., 2014. Fungal chitin dampens inflammation through IL-10 induction mediated by NOD2 and TLR9 activation. PLoS Pathog 10, e1004050. 10.1371/journal.ppat.1004050

Waheed, A.A., Rao, K.S., Gupta, P.D., 2000. Mechanism of dye binding in the protein assay using eosin dyes. Anal. Biochem. 287, 73–79. 10.1006/abio.2000.4793

Walsh, N.M., Wüthrich, M., Wang, H., Klein, B., Hull, C.M., 2017. Characterization of C-type lectins reveals an unexpectedly limited interaction between *Cryptococcus neoformans* spores and Dectin-1. PLoS ONE 12, e0173866. 10.1371/journal.pone.0173866

Wang, Z.A., Griffith, C.L., Skowyra, M.L., Salinas, N., Williams, M., Maier, E.J., Gish, S.R., Liu, H., Brent, M.R., Doering, T.L., 2014. *Cryptococcus neoformans* dual GDP-mannose transporters and their role in biology and virulence. Eukaryot. Cell 13, 832–842. 10.1128/ec.00054-14

Winski, C.J., Stuckey, P.V., Marrufo, A.M., Agyei, G., Ross, R.L., Urmi, T., Chapman, S., Santiago-Tirado, F.H., 2025. Lack of an atypical PDR transporter generates an immunogenic *Cryptococcus neoformans* strain that drives a dysregulated and lethal immune response in murine lungs. mBio 16, e01321–25. 10.1128/mbio.01321-25

