## supplement-figures for "Novel cell wall-associated genes that enable *Cryptococcus neoformans* to evade dectin-1-mediated innate immune recognition"

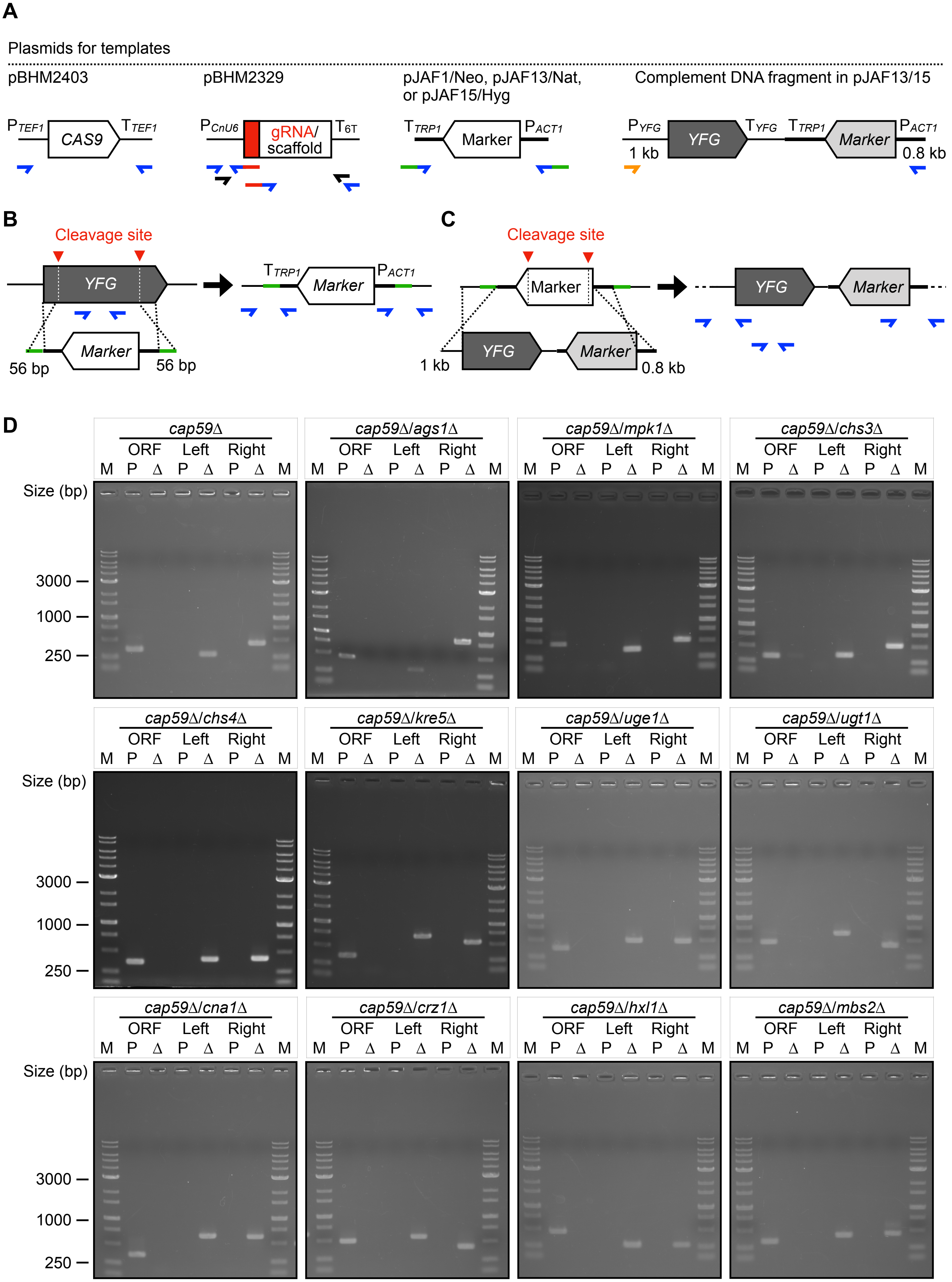


**Figure S1 continued**

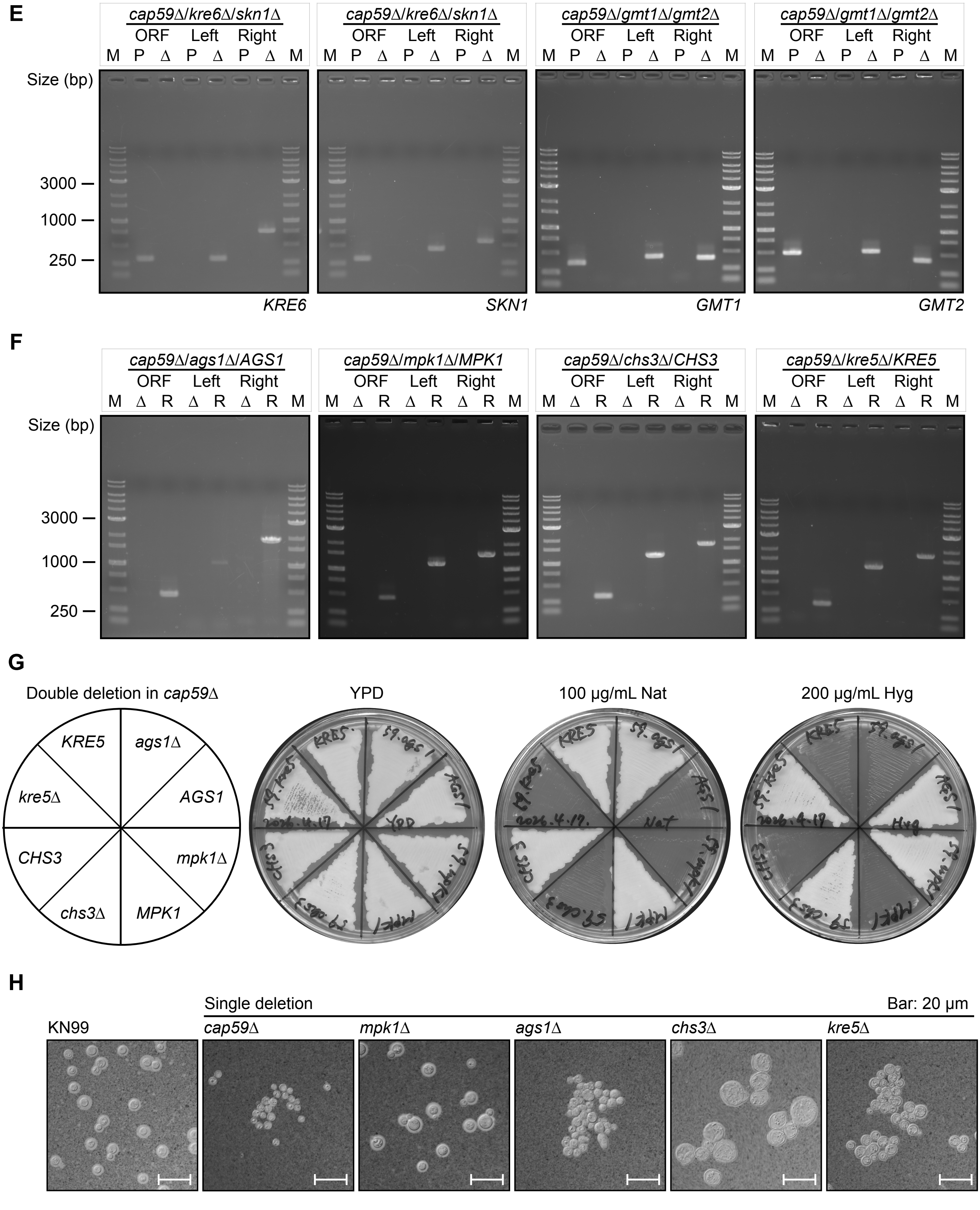
E
cap59Δ/kre6Δ/skn1Δ
cap59Δ/gmt1Δ/gmt2Δ
ORF Left Right
M P Δ P Δ P Δ M
Size (bp)
3000
1000
250
KRE6
SKN1
GMT1
GMT2
F
cap59Δ/ags1Δ/AGS1
cap59Δ/mpk1Δ/MPK1
cap59Δ/chs3Δ/CHS3
cap59Δ/kre5Δ/KRE5
M Δ R Δ R Δ R M
G
Double deletion in cap59Δ
YPD
100 μg/mL Nat
200 μg/mL Hyg
KRE5
ags1Δ
kre5Δ
AGS1
CHS3
mpk1Δ
chs3Δ
MPK1
H
Single deletion
Bar: 20 μm
KN99
cap59Δ
mpk1Δ
ags1Δ
chs3Δ
kre5Δ

**Figure S1**

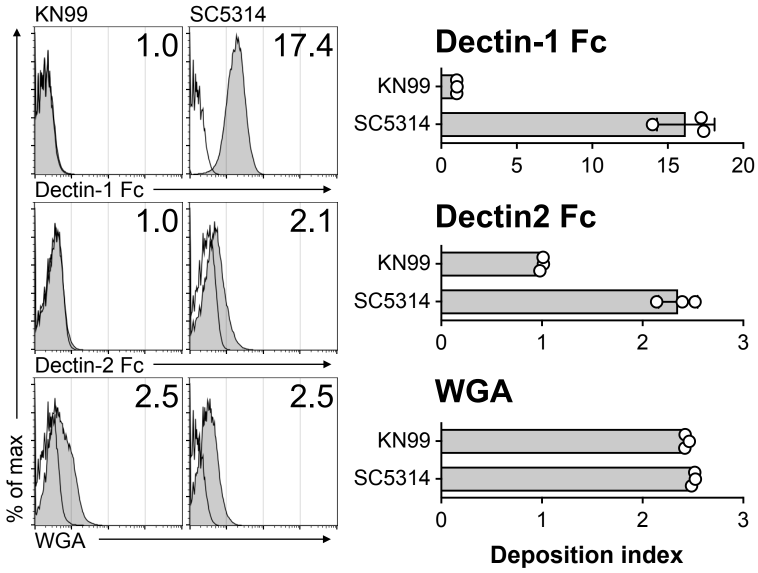
KN99
SC5314
1.0
17.4
Dectin-1 Fc
1.0
2.1
Dectin-2 Fc
2.5
2.5
% of max
WGA
Dectin-1 Fc
KN99
SC5314
0
5
10
15
20
Dectin2 Fc
KN99
SC5314
0
1
2
3
WGA
KN99
SC5314
0
1
2
3
Deposition index

**Figure S2**

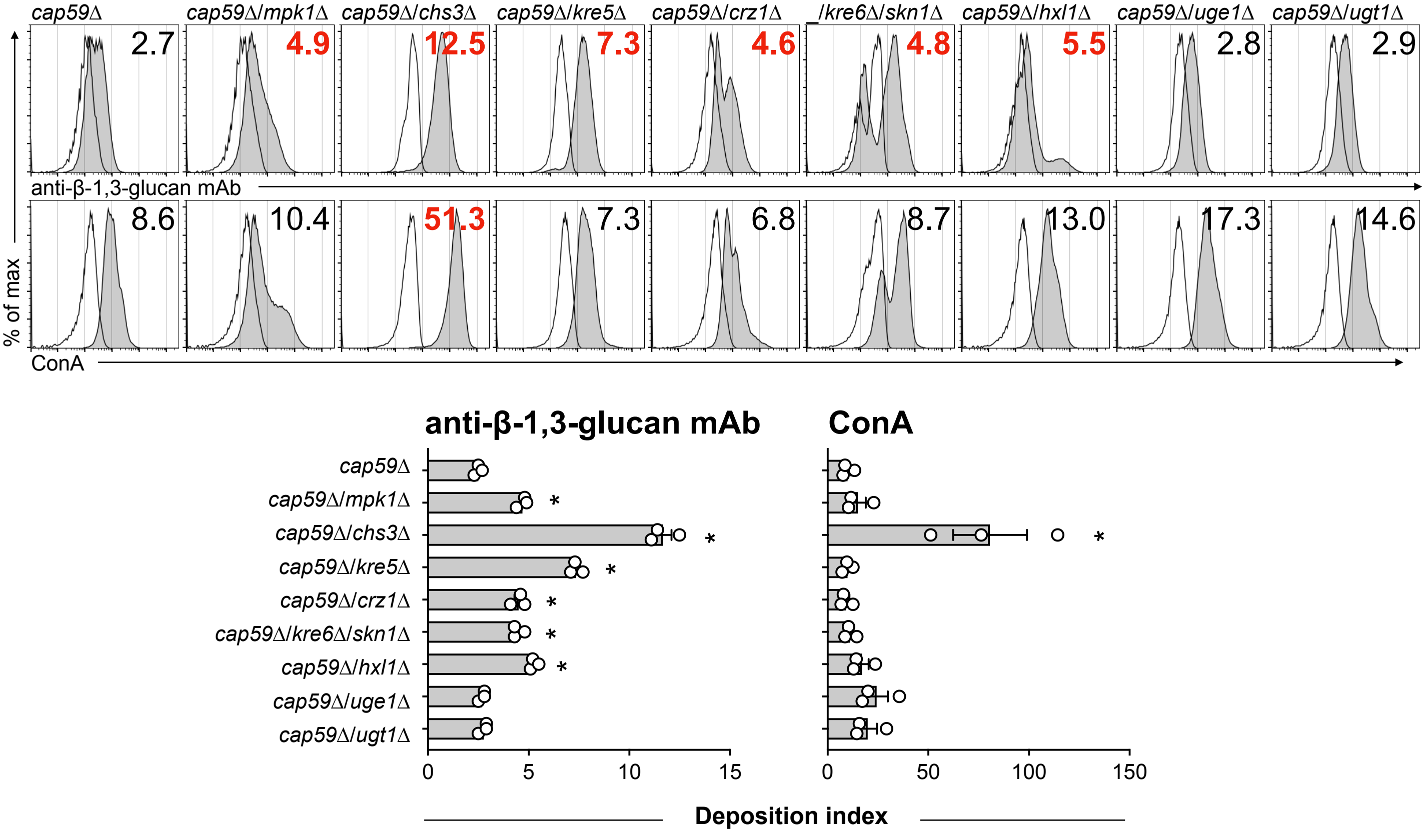
cap59Δ
cap59Δ/mpk1Δ
cap59Δ/chs3Δ
cap59Δ/kre5Δ
cap59Δ/crz1Δ
_/kre6Δ/skn1Δ
cap59Δ/hxl1Δ
cap59Δ/uge1Δ
cap59Δ/ugt1Δ
2.7
4.9
12.5
7.3
4.6
4.8
5.5
2.8
2.9
anti-β-1,3-glucan mAb
8.6
10.4
51.3
7.3
6.8
8.7
13.0
17.3
14.6
% of max
ConA
anti-β-1,3-glucan mAb
ConA
cap59Δ
cap59Δ/mpk1Δ
cap59Δ/chs3Δ
cap59Δ/kre5Δ
cap59Δ/crz1Δ
cap59Δ/kre6Δ/skn1Δ
cap59Δ/hxl1Δ
cap59Δ/uge1Δ
cap59Δ/ugt1Δ
0
5
10
15
0
50
100
150
Deposition index

**Figure S3**

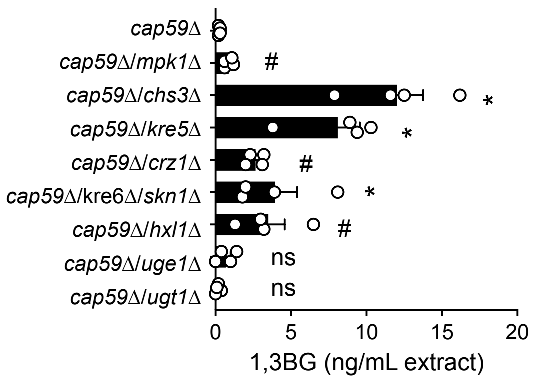
cap59Δ
cap59Δ/mpk1Δ
cap59Δ/chs3Δ
cap59Δ/kre5Δ
cap59Δ/crz1Δ
cap59Δ/kre6Δ/skn1Δ
cap59Δ/hxl1Δ
cap59Δ/uge1Δ
cap59Δ/ugt1Δ
#
*
*
#
*
#
ns
ns
0
5
10
15
20
1,3BG (ng/mL extract)

**Figure S4**

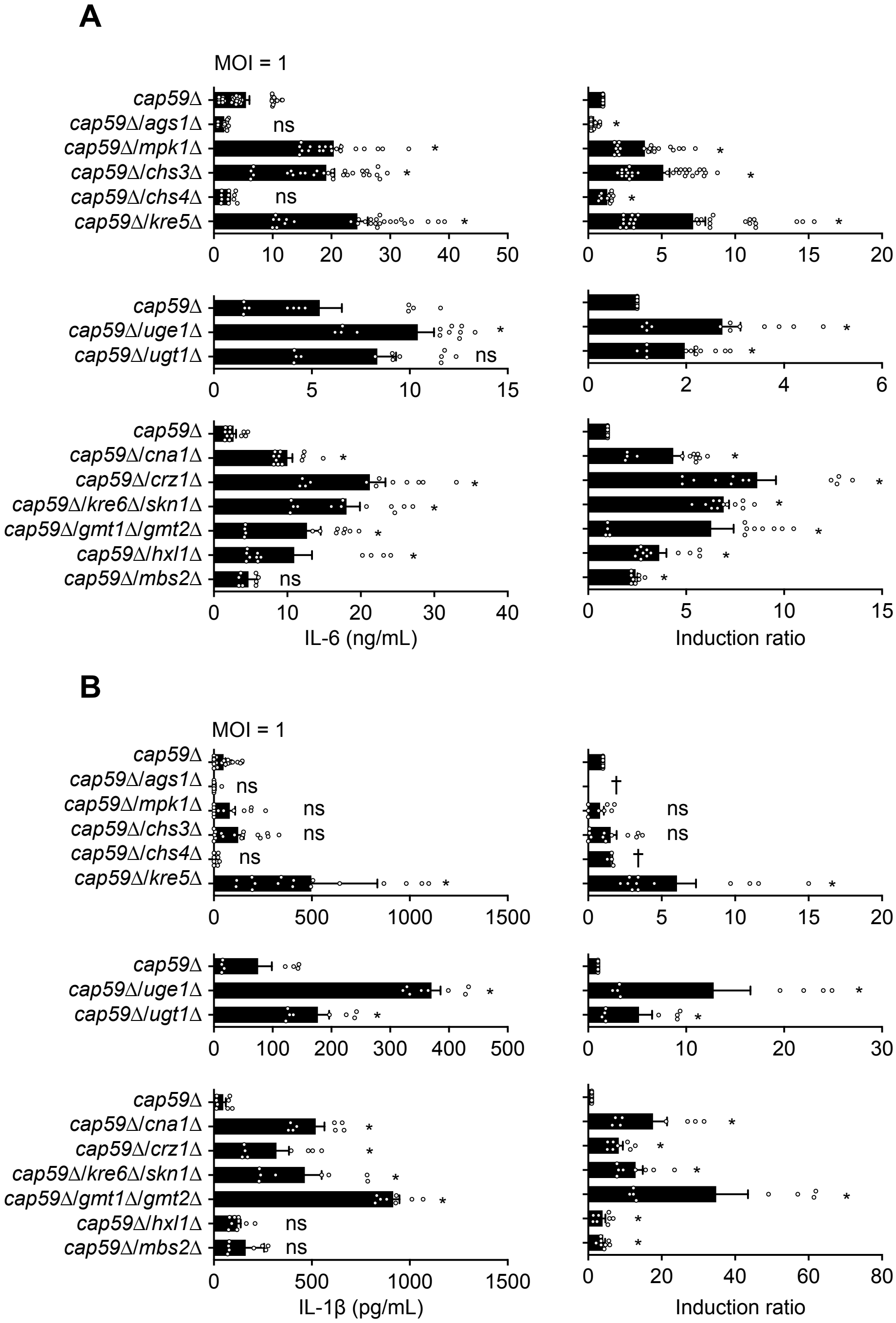
A
MOI = 1
cap59Δ
cap59Δ/ags1Δ
cap59Δ/mpk1Δ
cap59Δ/chs3Δ
cap59Δ/chs4Δ
cap59Δ/kre5Δ
cap59Δ/uge1Δ
cap59Δ/ugt1Δ
cap59Δ/cna1Δ
cap59Δ/crz1Δ
cap59Δ/kre6Δ/skn1Δ
cap59Δ/gmt1Δ/gmt2Δ
cap59Δ/hxl1Δ
cap59Δ/mbs2Δ
ns
IL-6 (ng/mL)
Induction ratio
B
MOI = 1
IL-1β (pg/mL)
Induction ratio
†

**Figure S5**

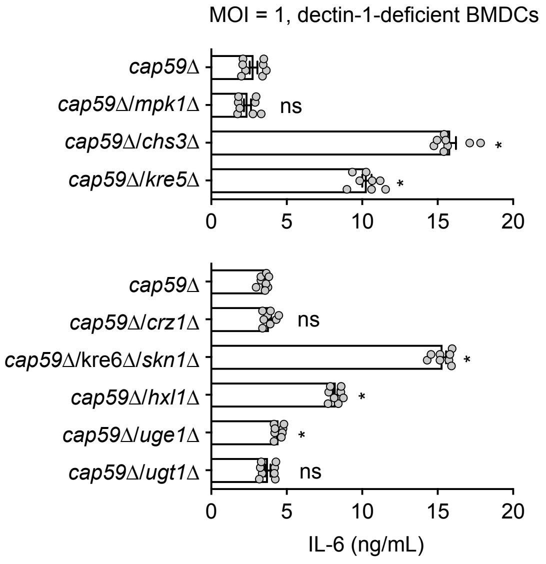

MOI = 1, dectin-1-deficient BMDCs
cap59Δ
cap59Δ/mpk1Δ
cap59Δ/chs3Δ
cap59Δ/kre5Δ
ns
*
*
0
5
10
15
20
cap59Δ
cap59Δ/crz1Δ
cap59Δ/kre6Δ/skn1Δ
cap59Δ/hxl1Δ
cap59Δ/uge1Δ
cap59Δ/ugt1Δ
ns
*
*
*
ns
0
5
10
15
20
IL-6 (ng/mL)


**Figure S6**

### 1,3BG masking-related genes in *C. neoformans*

| This study | Bloom et al., Nat commun, 2019 |
|---|---|
| YPD 30°C (Standard conditions) | YPD 37°C (Thermal stress) |
| • Protein kinase: *MPK1*<br>• Transcriptoinal factor: *CRZ1*, *HXL1*<br>• Cell wall synthesis (Chitosan): *CHS3*<br>• Cell wall synthesis (1,6BG): *KRE5*, *KRE6*/*SKN1*<br>• Cell wall synthesis (GXMGal): *UGE1*, *UGT1* | • mRNA decay (mRNA deadenylase): *CCR4* |
| Deletant phenotype | |
| • Compensatory cell wall rearrangement<br>• 1,3BG exposure and dectin-1 deposition<br>• Highly immunogenic to BMDCs (IL-6/IL-1β) | • Aberrant transcriptional reprogramming<br>• 1,3BG exposure and dectin-1 deposition |

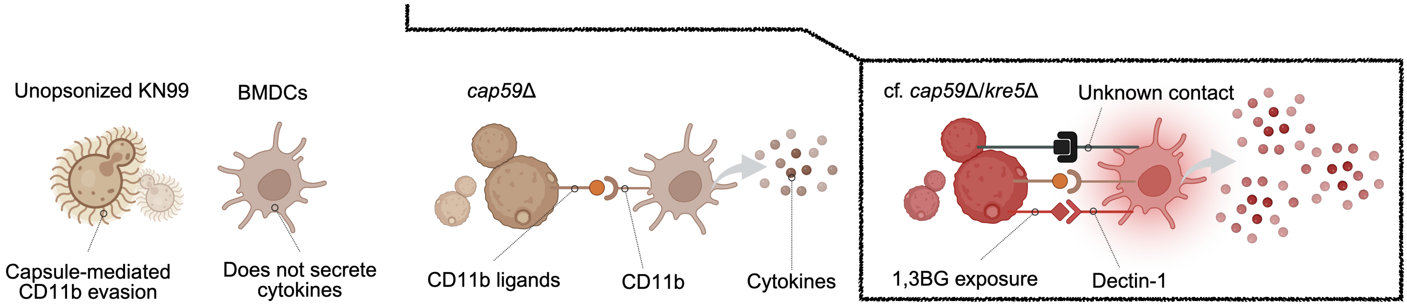


**Figure S7**
